# Quality control of large genome datasets using genome fingerprints

**DOI:** 10.1101/600254

**Authors:** Max Robinson, Arpita Joshi, Ansh Vidyarthi, Mary Maccoun, Sanjay Rangavajjhala, Gustavo Glusman

## Abstract

The 1000 Genomes Project (TGP) is a foundational resource which serves the biomedical community as a standard reference cohort for human genetic variation. There are now seven public versions of these genomes. The TGP Consortium produced the first by mapping its final data release against human reference sequence GRCh37, then “lifted over’’ these genomes to the improved reference sequence (GRCh38) when it was released, and remapped the original data to GRCh38 with two similar pipelines. As ‘best practice’ quality validation, the pipelines that generated these versions were benchmarked against the Genome In A Bottle Consortium’s ‘platinum quality’ genome (NA12878). The New York Genome Center recently released the results of independently resequencing the cohort at greater depth (30X), a phased version informed by the inclusion of related individuals, and independently remapped the original variant calls to GRCh38. We evaluated all seven versions using genome fingerprinting, which supports ultrafast genome comparison even across reference versions. We noted multiple issues including discrepancies in cohort membership, disagreement on the overall level of variation, evidence of substandard pipeline performance on specific genomes and in specific regions of the genome, cryptic relationships between individuals, inconsistent phasing, and annotation distortions caused by the history of the reference genome itself. We therefore recommend global quality assessment by rapid genome comparisons, using genome fingerprints and other metrics, alongside benchmarking as part of ‘best practice’ quality assessment of large genome datasets. Our observations also help inform the decision of which version to use, to support analyses by individual researchers.

## Introduction

Since its initial release, the Thousand Genomes Project (TGP)^1^ has served its intended purpose as the standard reference for human genetic variation in population structure analyses, genotype imputation efforts, association studies, evaluations of gene annotation, and efforts to improve the reference genome itself, among many other applications.^2^ Since its release in 2013, most analyses of the TGP cohort have relied on the ‘phase 3’ final data release version, which includes estimated variation in 2504 individuals mapped onto version GRCh37 (hg19) of the human reference genome. These individuals were sampled from 26 populations in 5 continent-level regions (Africa, East Asia, South Asia, Europe, and the Americas). Genotypes for all individuals were estimated based on a combination of low-coverage whole-genome sequencing, deep exome sequencing, and high-density single-nucleotide polymorphism (SNP) microarrays. The resulting variant calls included phased biallelic and multiallelic SNPs, indels and structural variants, with high power to detect variants with alleles of at least 5% frequency (75-95% depending on population)^1^. For the purpose of comparing variation in the same individuals relative to different reference sequences, we hereafter refer to this phase 3 dataset as **TGP37**.

Not long after the initial release of TGP37, a new and much improved version^3^ of the human reference sequence, GRCh38 (hg38), was released, prompting efforts to express this standard set of variation data relative to this new reference. In 2015, the subset of TGP37 variants with reference SNP ID numbers (rsids)^4^ were translated (via liftOver) to GRCh38 coordinates, yielding a version of the variation reference set we will call **TGP38L** (‘L’ for ‘lifted’). In 2017, the raw genomic sequence reads were mapped^5^ onto GRCh38 to support ‘native’ variant calling.^6^ Two versions of these variant calls have been released to date via the International Genome Sample Resource (IGSR).^7^ The first, released in late 2018, is restricted to biallelic single-nucleotide variants (SNVs); we call this version **TGP38S** (‘S’ for ‘SNVs’). The second, released in early 2019, includes both biallelic SNVs and indels; we refer to this extended dataset as **TGP38X** (‘X’ for ‘extended’). Also in early 2019, a new set of high-coverage whole-genome sequences were produced by the New York Genome Center (NYGC) and released via the European Nucleotide Archive (accession: PRJEB31736) and IGSR; we refer to this set as **TGP38H** (‘H’ for ‘high coverage’).^8^ This new set was later (May 2020) enhanced by the addition of the genomes of 698 related individuals, leading to improved phasing; we refer to this improved set (limited to just the original 2,504 individuals in TGP) as **TGP38N** (‘N’ for ‘new’).

The NYGC also remapped the original variant calls in TGP37 using CrossMap^9^, yielding the version we call **TGP38C** (‘C’ for ‘crossmap’). We anticipate that one of these versions will soon become the new standard reference version of the human genetic variation in the TGP cohort. As such, it is important to evaluate the quality of each of these genome sets, to enable researchers to choose the most appropriate version to support their specific research questions.

Despite the intent to include only unrelated individuals in the TGP cohort,^10^ a number of close and more distant relationships exist within TGP37, as reported by us and others.^11^ TGP37 is supplemented with a small set of 31 related individuals, which we call **TGP37r** (‘r’ for ‘related’). Likewise, a set of 150 related samples accompanies TGP38S and TGP38X - we refer to these in turn as **TGP38Sr** and **TGP38Xr**. Finally, we refer to the aforementioned 698 genomes accompanying TGP38N as **TGP38Nr**.

Given the importance of this high-impact reference set of human variation, validating the quality of each release is a crucial concern. TGP37 data were evaluated by comparing variant calls for one individual per population to calls based on 30x PCR-free sequence data, and by comparison to 47x sequence using Complete Genomics technology.^1^ TGP38S data quality were evaluated prior to release by comparing variant calls for the NA12878 individual to ‘truth set’ calls from the Genome In A Bottle (GIAB) Consortium.^6, 12–14^ At least two independent studies of TGP data quality have also been reported. A study of the array genotypes for 2318 TGP samples found the data to be in general of high quality, though some discrepancies were found between reported and inferred sex or in degree of relationship for some samples.^15^ More recently, a study found that phasing and imputation for rare variants are unreliable, based on comparison to haplotypes of 28 individuals, experimentally phased using a different sequencing technology.^16^

The availability of multiple versions of the same dataset enables a different type of quality control: cross-comparison among the versions, which supports identification of version-specific results and data processing failures throughout the cohort, the majority of which is not assessed by benchmarking of results against a small subset of high-quality genomes. Cross-comparisons can be performed very efficiently using reduced genome representations, including genome fingerprints.^11^ Here, we use genome fingerprints and some additional statistics to compare the seven versions of the TGP (TGP37, TGP38L, TGP38C, TGP38S, TGP38X, TGP38H, and TGP38N) and their associated related samples (TGP37r, TGP38Sr, TGP38Xr, and TGP38Nr), in terms of (a) the set of genomes analyzed, (b) known and cryptic relatedness within each cohort, (c) patterns of SNV and genotype concordance between versions, and (d) phasing concordance. We then seek explanations for the results of these comparisons with a limited set of rapid follow-up studies.

## Materials and Methods

### Datasets

We obtained seven versions of the 1000 genomes dataset, phase 3, from IGSR.^7^

1. **TGP37**: Variant calls relative to the GRCh37 (hg19) version of the human genome reference (N=2504).
2. **TGP38L**: Variant calls for the same set of genomes, “lifted over” to the GRCh38 (hg38) version of the human reference using dbSNP v. 149 (N=2504). This version was later withdrawn (in March 2021).
3. **TGP38C**: Variant calls for the same set of genomes, “lifted over” to the GRCh38 (hg38) version of the human reference using CrossMap (N=2504).
4. **TGP38S**: A set of integrated phased biallelic SNV calls, directly called against the GRCh38 (hg38) version of the human reference (N=2548).
5. **TGP38X**: An extended set of integrated phased biallelic SNV and indel calls, directly called against the GRCh38 (hg38) version of the human reference (N=2548).
6. **TGP38H**: A high-coverage (30x) set of recalibrated SNV and indel calls, produced by the New York Genome Center, directly called against GRCh38 with alternative sequences and decoys (N=2504) - European Network Archive Project <u>PRJEB31736</u>.
7. **TGP38N**: A high-coverage (30x) set of recalibrated SNP and indel calls, produced by the New York Genome Center, directly called against GRCh38 with alternative sequences and decoys (N=2504), and phased in combination with 698 related individuals - European Network Archive Study ERP114329, Project <u>PRJEB31736</u>. We downloaded VCF files including all 3202 genomes, and split them into 2504 main, and 698 relateds, for further analysis.

We also obtained four sets of genomes of samples related to those in the main TGP dataset.

1. **TGP37r**: Integrated phased biallelic SNV and indel calls, relative to GRCh37 (N=31).
2. **TGP38Sr**: Integrated phased biallelic SNV calls, relative to GRCh38 (N=150).
3. **TGP38Xr**: Integrated phased biallelic SNV and indel calls, relative to GRCh38 (N=150).
4. **TGP38Nr**: Integrated phased SNV and indel calls, relative to GRCh38 (N=698) - European Network Archive Study ERP120144, Project <u>PRJEB36890</u>.

### Variant normalization

We normalized the variants in TGP38C and TGP38N using the `bcftools norm -m+` command^17^.

### Whole-genome fingerprinting

We computed fingerprints for all genomes in all sets as described,^11^ with L=200. Unless otherwise specified, all genome fingerprints include only biallelic autosomal SNVs. This computation does not include the pseudoautosomal regions (PARs) of chromosomes X and Y.

### Chromosome fingerprinting

To compute single-chromosome fingerprints, we restricted SNV pair collection to each chromosome and normalized the single-chromosome raw fingerprints separately, yielding single-chromosome normalized fingerprints. Other than restricting the range to the individual chromosome, the procedure is identical to that used for computing whole-genome fingerprints. We applied this procedure also to the PARs.

### Other metrics

We computed the *SNV count* of an individual as the number of biallelic SNVs observed in their genome in either heterozygous state or homozygous for the alternate allele. We computed the *genotype concordance* of an individual between two datasets as the number of biallelic SNVs (with same coordinates, REF and ALT alleles in the two datasets) in which the individual is heterozygous in both datasets (ignoring phasing of heterozygous sites) or homozygous alternate allele in both datasets, divided by the total number of biallelic SNVs in which the individual was not homozygous reference in both datasets.

### Evaluation of sex chromosome coverage

We downloaded CRAM index files for all samples from the STU population (*.alt_bwamem_GRCh38DH.20150718.STU.low_coverage.cram.crai) and used these to estimate total coverage in chrX and chrY (and hence estimate chrX and chrY copies) by normalizing to the observed coverage in autosomes. We similarly obtained and analyzed CRAM index files for the 698 samples in TGP38Nr.

### Identification of genomic regions discrepant between TGP versions

We aligned the VCF files for each pair of TGP versions, to identify variants shared between the two versions (same chromosome, position, reference and alternate alleles) and to enumerate variants unique to one version (i.e., absent from the other). When encountering variants in which all individuals were reported as homozygous reference (i.e., zero counts of the alternate allele, AC==0), we considered them as absent. We then identified segments enriched in (or depleted of) unique variants, using this procedure: for each pair of consecutive shared variants, we counted how many unique variants are present between them in each of the two TGP versions being compared, and retained for subsequent analysis segments at least 1 kb long. We identified a subset of these segments as significant, based on p-values returned by the corresponding cumulative distribution function of the Poisson distribution, modeled using the length of the segment and the number of unique variants in it (separately for each TGP version being compared). The Poisson distribution is used to estimate the number of events in a given time or space interval. It uses a shaping parameter *lambda* (λ) which denotes the expected number of events in a unit of time or space. We use the genome wide density of variants to estimate λ for a given segment of length *l*.

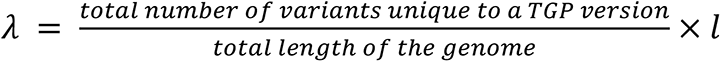

The p-value to determine the extremity of occurrence of *k* variants in a segment is then evaluated as:

*if*(*k* > *λ*)

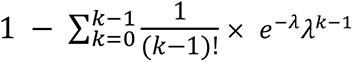

*else:*

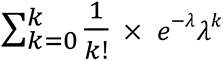

Using Bonferroni correction, we considered significant any segment with *p-value* ≤ 0.05/2*N*, where N=104,572 is the total number of segments being evaluated in all pairwise comparisons (two p-values are computed per segment). We then used bedtools^18^ to subtract from the list of ‘unique segments’ those that overlap with large genomic gaps or satellite sequences (yielding a list of 95,843 segments), and finally to merge them into a set of 12,307 non-overlapping regions.

### Identification and classification of discrepant alternate alleles

We compared all six TGP versions on GRCh38 (TGP38L, TGP38S, TGP38X, TGP38H and TGP38N) to identify autosomal variants in which the alternate allele (ALT) differs between TGP versions. To maximize interpretability, we restricted this analysis to the 2503 individuals represented in all TGP versions. We tabulated the reported ALT alleles and their allele counts (AC) in each TGP version, and classified the ALT-discrepant variants between TGP38L and TGP38X into three non-overlapping classes: (1) *Multiallelic*, including variants with more than two ALT alleles in any TGP version; (2) *Indel-related*, including variants in which the REF allele or any ALT allele has length different from 1, or one of the alleles is a ‘star allele’ (the * character), in any TGP version; and (3) *Biallelic SNV*, including all remaining ALT-discrepant biallelic single-nucleotide variants not obviously associated with indels in any version of TGP. Sites that are both multiallelic and indel-related are counted among the multiallelic.

### Phasing-aware genome fingerprints

We modified the fingerprint-computation method to capture local phasing information by changing the third step of raw fingerprint computation.^11^ Specifically, when joining the keys of consecutive SNVs, we reverse the key of the second SNV (i.e., the ALT allele is assigned to the first phased chromosome and the REF allele is assigned to the second chromosome) if only one of the two SNVs matches the genotype pattern “1|0”.

We computed phasing-aware genome fingerprints for all TGP versions except for TGP38H, since this version is not phased. Finally, we compared the datasets in pairwise fashion using standard and phasing-aware fingerprints.

## Results

### Overview of datasets and quality assessment strategies

We demonstrate the application of genome fingerprints^11^ for rapid cross-comparison of large genome datasets on the seven reported versions of the 1000 Genomes phase 3 cohort (Table 1 and Figure 1), which included the variant set as originally released, mapped to GRCh37 (TGP37); the same variants lifted over to GRCh38 (TGP38L and TGP38C); a reanalysis of the same raw sequence reads directly mapped onto GRCh38, first limited to biallelic SNVs (TGP38S) and then also including biallelic indels (TGP38X); as well as a recent independent high-coverage resequencing (TGP38H and TGP38N) of the original cell lines (sequenced to a nominal target depth of 30X coverage; see Methods). We also evaluate smaller sets of related individuals (Figure 2).

**Figure 1.**
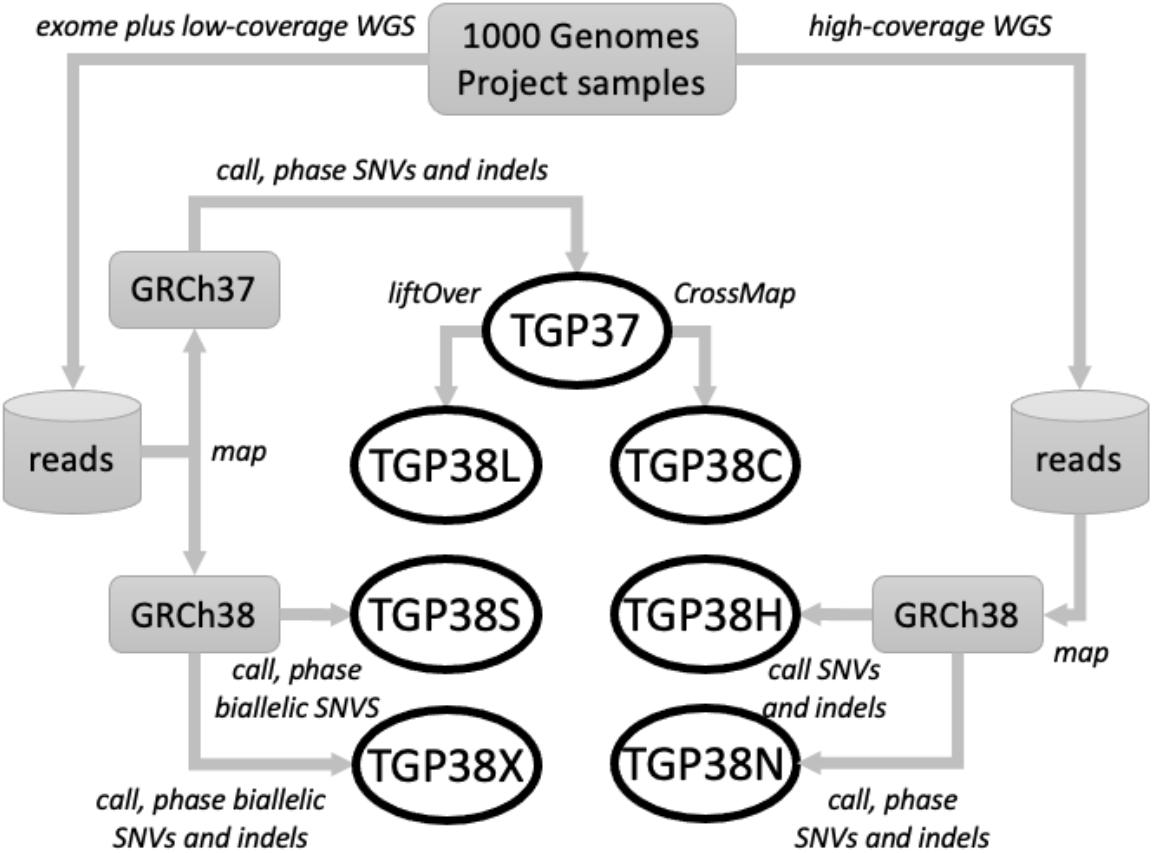
Overview of the seven available versions of sequence variations within the TGP phase 3 cohort. Phase 3 of the 1000 Genomes mapped exome sequences and low-coverage WGS onto GRCh37 to identify sequence variations, yielding version TGP37. Three approaches have since been used to represent the human genomic sequence variation against the GRCh38 reference sequence: liftOver of variants called against GRCh37 (yielding version TGP38L) and similarly using CrossMap (yielding version TGP38C); remapping the individual reads from the original data directly against GRCh38, followed by integrated variant calling resulting in only SNV calls (TGP38S) or both SNV and indel calls (TGP38X); and independent whole genome resequencing of new reads, for these 2504 sequences at the modern standard sequencing depth of 30X, (yielding versions TGP38H and TGP38N, which is also phased).

**Figure 2.**
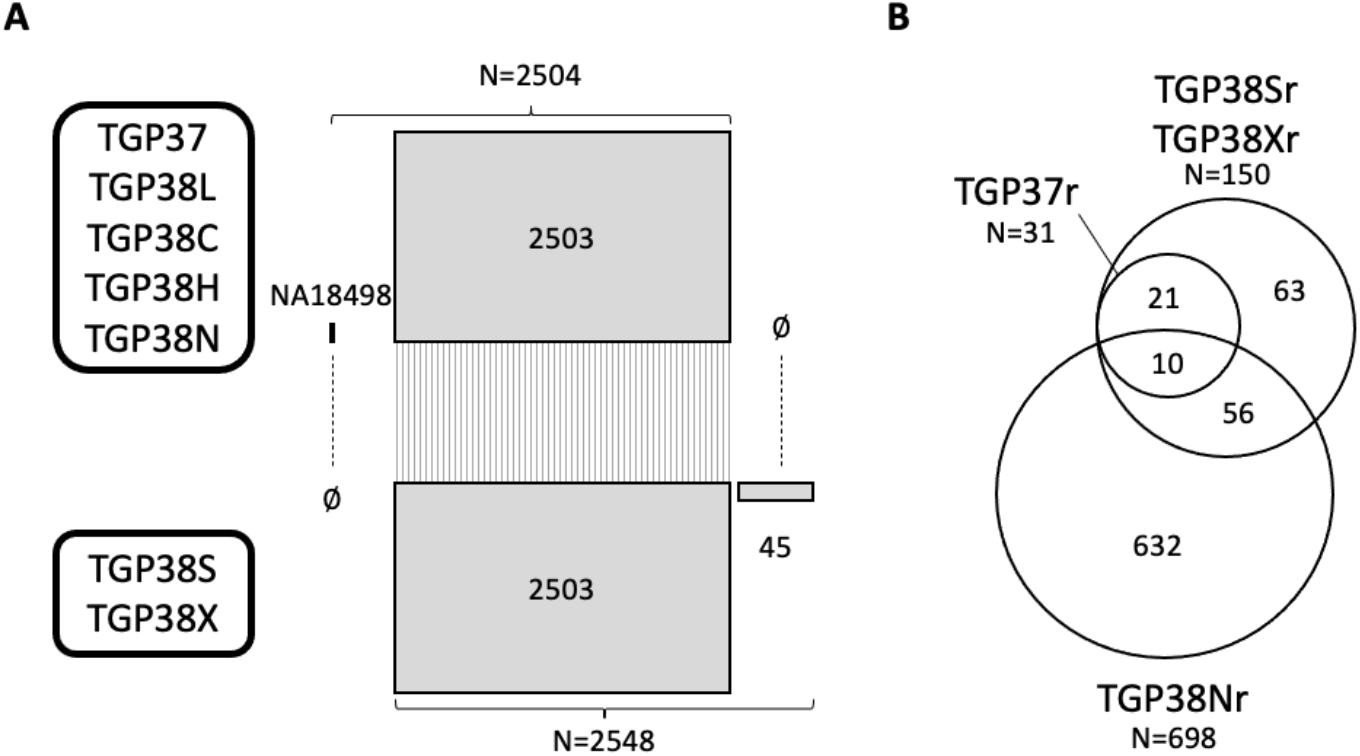
Comparison of cohort structure. We counted sequence identifiers in each TGP set, and came to the following conclusions. **A.** While TGP38L, TGP38H and TGP38N include the same 2504 genome identifiers as TGP37, as expected, TGP38X and TGP38S include 2548 identifiers, only 2503 of which are in common with TGP37. NA18498 is absent from versions TGP38S and TGP38X, which also include 45 identifiers not in TGP37. **B.** The original set of 31 supplemental ‘related samples’ (TGP37r) has been expanded to 150 in the versions TGP38Sr and TGP38Xr. TGP38Nr includes 698 genomes with partial overlap with the other sets.

**Table 1.** Overview of datasets.

| TGP version | TGP37 | TGP38L | TGP38C | TGP38S | TGP38X | TGP38H | TGP38N |
| --- | --- | --- | --- | --- | --- | --- | --- |
| Data | exome plus low-coverage WGS |  |  |  |  | 30x WGS |  |
| Method | map and call | liftOver | CrossMap | map and call |  |  |  |
| Reference version | GRCh37 | GRCh38 |  |  |  |  |  |
| Reference file | hs37d5 |  | not stated | GRCh38_full_analysis_set_plus_decoy_hla |  |  |  |
| Individuals | 2,504 |  |  | 2,503 + 45 |  | 2,504 |  |
| Related individuals | 31 | NA |  | 150 |  | NA | 698 |
| File format | VCFv4.1 |  |  | VCFv4.3 |  | VCFv4.2 | VCFv4.1 |
| ##source header | 1000GenomesPhase3Pipeline |  | not stated | IGSRpipeline |  | not stated |  |
| ##fileDate header date | 20140730 | 20150218 | not stated | 31052018 | 26022019 | not stated |  |
| Autosomes | 1-22 |  | chr1-chr22 | chr1-chr22 | 1-22 | chr1-chr22 |  |
| Sex chromosomes | X, Y |  | chrX, chrY | chrX | X | chrX, chrY | chrX |
| Mitochondrial chromosome | MT | - |  |  |  | chrM | - |
| SNVs included | all |  |  | only biallelic |  | all |  |
| Indels included | all |  |  | none | only biallelic | all |  |
| Multiallelic representation | single-line |  | multi-line | NA |  | single-line | multi-line |
| Non-singleton clusters | 367,892 | 361,831 | 747,172 | 0 | 712,705 | 2,783,314 | 2,696,557 |
| Clusters with multi-line SNVs | 1,940 | 267 | 259,159 | 0 | 20 | 0 | 593,152 |
| Variation lines | 81,271,745 | 81,045,153 | 81,184,416 | 73,159,510 | 78,122,255 | 120,046,355 | 66,616,026 |
| Biallelic SNVs | 77,805,856 | 77,639,718 | 77,661,355 | 73,159,510 | 72,458,679 | 104,726,921 | 59,129,084 |
| Variants with rsids | 48.3% | 100% | 99.9% | 0% |  |  |  |

| Phased | yes |  |  |  |  | no | yes |
| --- | --- | --- | --- | --- | --- | --- | --- |
| **Dataset**<br>**download size**<br>**(gigabytes)** | 15.8 | 16.2 | 16.3 | 12.9 | 14.2 | 1241.4 | 32.3 |

Beyond the method by which they were produced, the seven versions of the TGP dataset differ in several ways (Table 1). The number of individuals included in each differs. All versions use the Variant Call Format (VCF) to represent the data, but they use different versions of VCF, represent chromosomes in different ways, include varying subsets of the sex and mitochondrial chromosomes, and include different combinations of SNVs, indels, biallelic or multiallelic variants. The number of variants reported for each individual can vary significantly. Two of the versions (TGP38C and TGP38N) represent multiallelic variants in multiple lines, a format that may be unexpected to many researchers and may confuse analysis tools, requiring variant normalization for proper analysis. The various versions also differ in their level of inclusion of variant identifiers (rsids), and whether variants are phased or not. On a very practical level, they also differ substantially in the total size of the dataset to be downloaded for local analysis.

We used genome fingerprints as well as simple summary statistics to compare the SNVs reported in these genomes. Based on these analyses, we identified a number of discrepancies and quality issues, including an unexpectedly missing individual (Figure 2), cryptic relations, and a set of genomes with significantly fewer SNV counts (Figures 3 and 4). We further identified regions of the genome with excess variation, or significantly lacking variation, in some TGP versions (Figure 5) and discrepancies in the identity of the alternate allele (ALT), sometimes at high population frequency (Figure 6). Finally, we observed that phasing is very inconsistent between the TGP versions (Figure 7).

**Figure 3.**
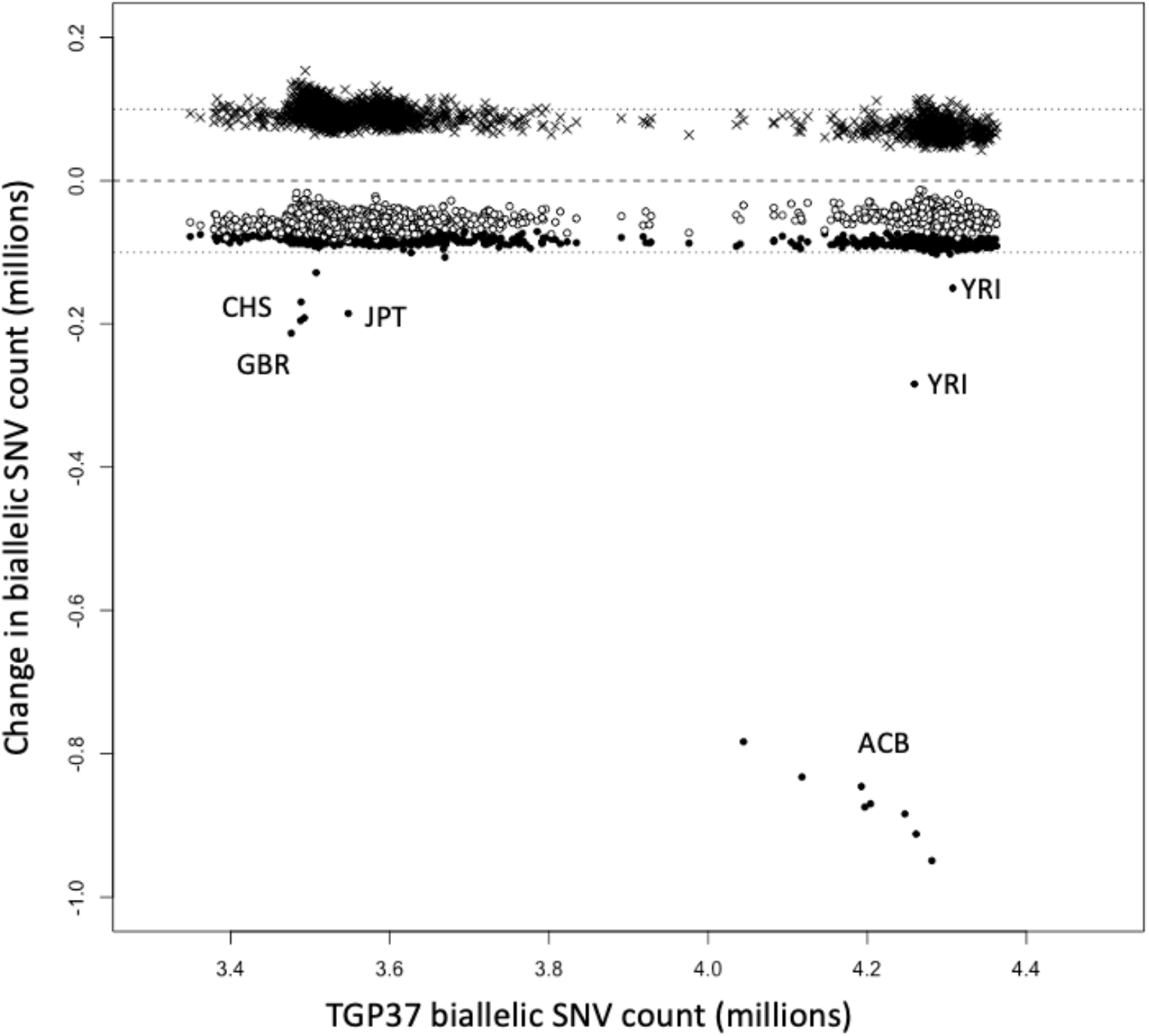
SNV count comparisons. Change in biallelic SNV counts (in millions) relative to TGP37, as observed in TGP38H (crosses), TGP38N (open circles) and TGP38X (filled circles). TGP38X and TGP38N universally report fewer SNVs than TGP37; 16 genomes are visibly exceptional in TGP38X. In particular, eight individuals from the ACB population have significantly fewer SNVs in TGP38X relative to TGP37. TGP38H universally reports more SNVs than TGP37, but without obvious outliers. Dashed line: no SNV count change. Dotted lines: addition (or loss) of 100,000 SNVs.

**Figure 4.**
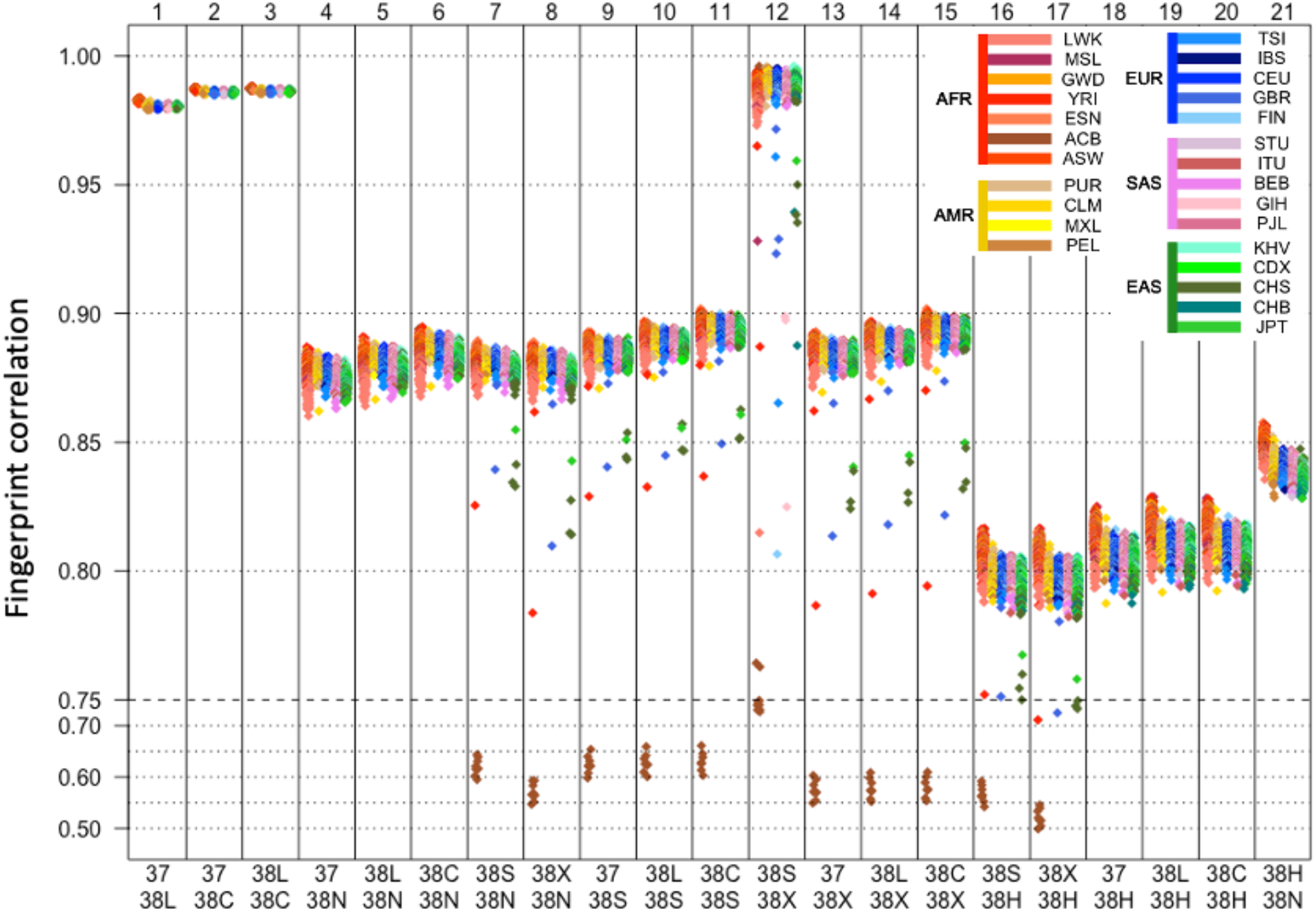
Distributions of fingerprint self-correlations in pairwise dataset comparisons. We compared fingerprints across TGP versions for each individual (color-coded by population), resulting in a distribution of Spearman’s ⍴ (y-axis) for each pair of versions (columns, labeled with numbers above, and with the versions being compared, below). Self-correlations of fingerprints for the same individual are expected to be high (⍴ > 0.75) irrespective of genome version or analysis pipeline^11^; lower self-correlations across versions show some genomes are unrecognizable as the same individual. Low outliers (⍴ < 0.75), are shown on a compressed scale under the dashed line.

**Figure 5.**
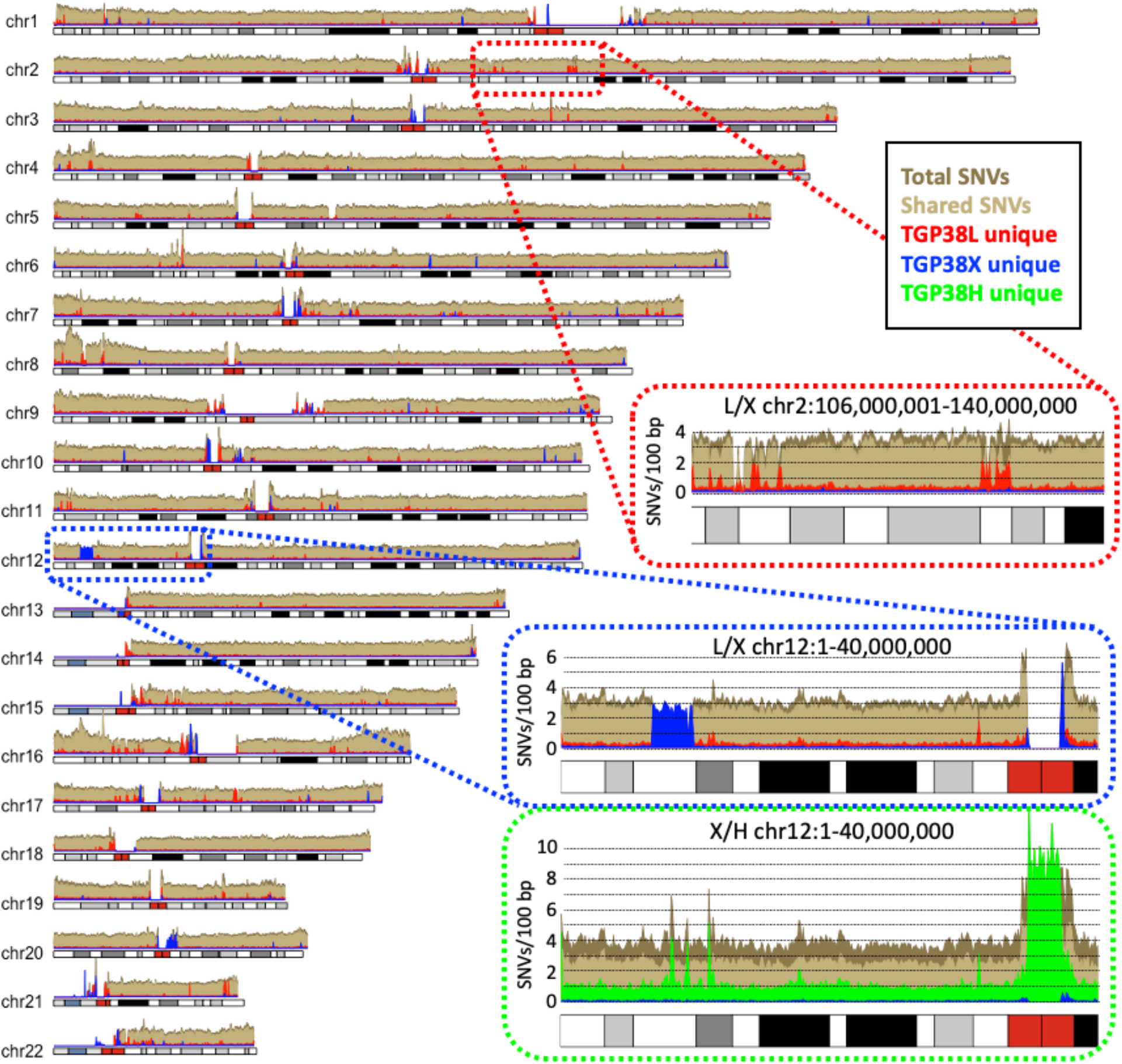
Density map of shared and unique variants along the chromosomes. The main plot shows the comparison between TGP38L and TGP38X, with shared variants in light brown, variants unique to TGP38L in red, those unique to TGP38X in blue, and the union of all variants in dark brown. Top inset (red outline): detail of a 36 Mb region on chr2 showing patches enriched in variants unique to TGP38L (absent from TGP38X). Middle inset (blue outline): detail of the first 40 Mb of chr12 showing a segment with only variants unique to TGP38X and no variation at all in TGP38L. Neither version has variants in the centromeric region. Lower inset (green outline): the same chr12 region when comparing TGP38X and TGP38H, highlighting the overall higher density of TGP38H-unique variants (in green), and the excess variation unique to TGP38H in centromeric regions.

**Figure 6.**
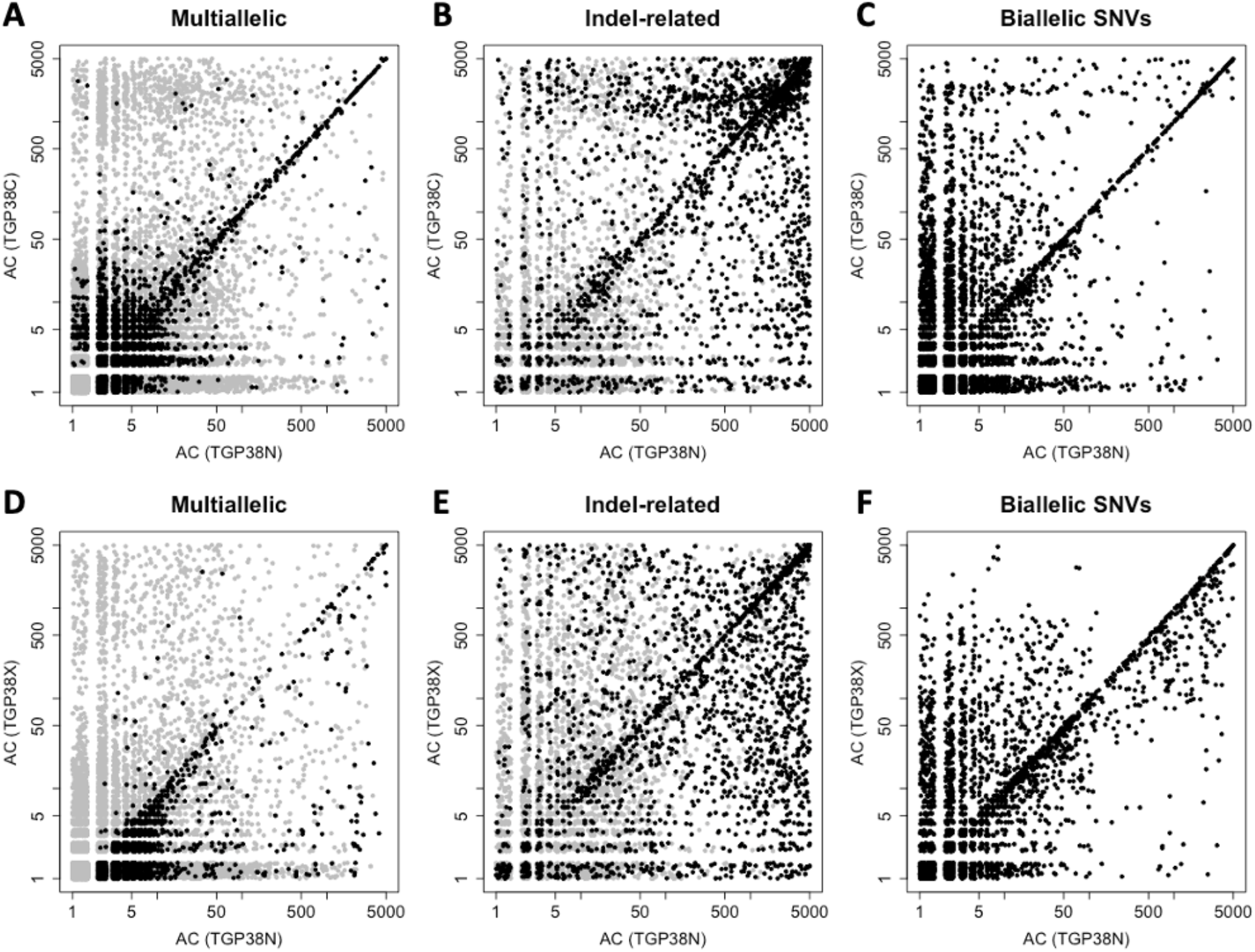
ALT-discrepant sites span the frequency range. Allele counts (AC) in log scale, jittered horizontally and vertically to denote density, for ALT-discrepant sites between TGP38N and the TGP version on the y-axis (TGP38C, panels **A**, **B** and **C**; TGP38X, panels **D**, **E** and **F**). **A** and **D**: Sites that are multiallelic in either version being compared (black) or in any of the other versions (gray). **B** and **E**: Indel-related sites in either version being compared (black) or in any of the other versions (gray). **C** and **F**: Biallelic SNVs not affected by indels.

**Figure 7.**
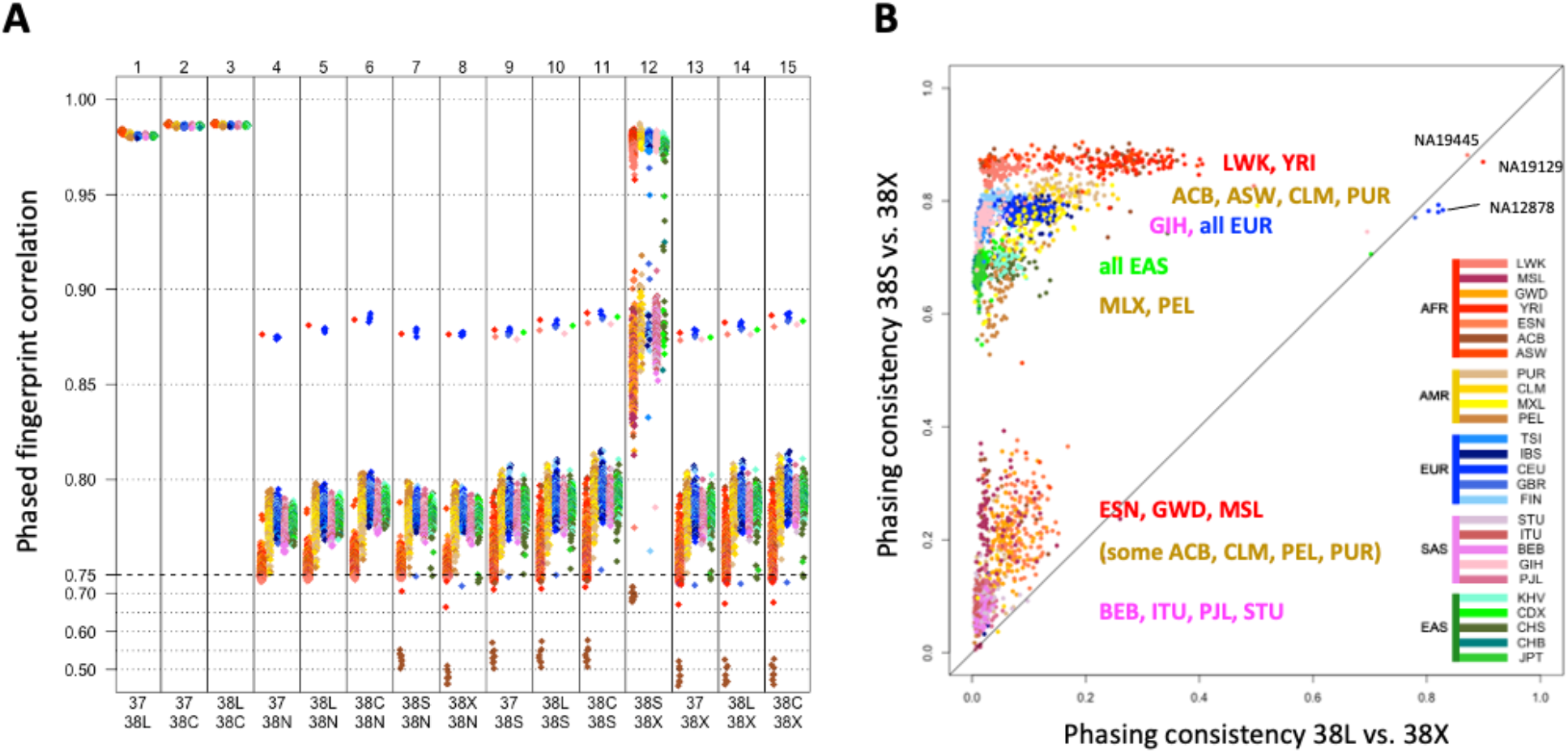
Consistency of phasing between TGP versions. **A.** We compared phase-aware fingerprints across TGP versions for each individual (color-coded by population), resulting in a distribution of Spearman’s ⍴ (y-axis) for each pair of versions (columns, labeled with numbers above, and with the versions being compared, below). Low outliers (⍴ < 0.75), are shown on a compressed scale under the dashed line. **B.** Using a phasing accuracy metric to compare TGP38L vs. TGP38X (x-axis) and TGP38S vs. TGP38X (y-axis) confirms the low phasing concordance for most genomes, and the differential phasing concordance by population, as observed in panel A, column 12.

### Cohort composition: metadata vs. genome comparison

We compared the identifiers of the genomes included in the seven TGP versions and observed that there are differences in cohort membership (Figure 2A), with four versions (TGP38L, TGP38C, TGP38H, and TGP38N) retaining the cohort used in TGP37, but the two versions directly mapping the original data to GRCh38 (TGP38S, TGP38X) omitting genome NA18498 (Yoruban in Ibadan, Nigeria, YRI), and including 45 additional genomes. The sets of related genomes also differ (Figure 2B), with 119 additional genomes (TGP38Sr, TGP38Xr) directly mapped to GRCh38 that were not in the original cohort (TGP37r), and 698 genomes in TGP38Nr, partially overlapping TGP37r and TGP38Xr. We found no overlap between the additional 45 genomes in TGP38X, and the 698 related genomes in TGP38Nr.

According to the metadata in the IGSR data portal,^19^ the omitted genome NA18498 should have been present in TGP38X. We used genome fingerprint comparisons to evaluate whether NA18498 might have been mislabeled, but found this not to be the case: the highest fingerprint correlation between NA18498 (from TGP38L) and any individual from TGP38X is 0.316 (to HG03108, Esan in Nigeria, ESN), which also exceeds correlation with any genome among the 150 supplemental individuals in TGP38Xr. These values are well below the 0.75 minimal correlation expected for versions of the same individual,^11^ confirming that NA18498 was not included in TGP38X or TGP38Xr under a different identifier. Of the 45 additional individuals in TGP38X, most (75%) seem to be related to individuals in the TGP37 cohort, some with fingerprint correlations consistent with second-degree relationships (> 0.45, Table S1), suggesting these genomes should not be included as part of the main TGP cohort.

Fingerprint-based comparisons of the 2503 individuals shared between TGP37 and TGP38X or TGP38S (Figure 2A) confirms a one-to-one relationship: for each individual in TGP37 (excluding NA18498, discussed above), the highest fingerprint correlation observed was to the TGP38X/TGP38S individual with the same identifier. For all but eight, the correlation is well above 0.75, as expected; the remaining eight individuals (‘strongly affected’, Table 2), all from the African Caribbean in Barbados (ACB) population, have between-set correlations that would be erroneously consistent with first-degree relationships.^11^ An additional eight individuals (‘mildly affected’, Table 2) have between-set fingerprint correlations ranging from 0.787 to 0.865, indicating the same individual (> 0.75) but much lower than observed for the other between-set self-comparisons (0.885 +/- 0.0028).

**Table 2.** Observed statistics for the outlier individuals most affected by dataset recomputation from TGP37 to TGP38X, in comparison to the ‘platinum’ NA12878 genome, the 89 ACB individuals unaffected by this bioinformatic difference, and the 2487 similarly unaffected individuals in the entire cohort. Genotype concordance is between TGP38L and TGP38X, as TGP37 and TGP38X are not on the same reference.

| Section | ID | Population code | Fingerprint correlation | SNV count (millions) |  | SNVs lost (%) | Genotype concordance |
| --- | --- | --- | --- | --- | --- | --- | --- |
|  |  |  |  | TGP37 | TGP38X |  |  |
| **missing** | NA18498 | YRI | NA | 4.28 | 0.00 | 100.00 | NA |
| **strongly affected** | HG02325 | ACB | 0.550 | 4.28 | 3.33 | 22.18 | 0.578 |
|  | HG02442 | ACB | 0.554 | 4.26 | 3.35 | 21.40 | 0.587 |
|  | HG02433 | ACB | 0.570 | 4.25 | 3.36 | 20.82 | 0.600 |
|  | HG01989 | ACB | 0.571 | 4.20 | 3.33 | 20.69 | 0.602 |
|  | HG02445 | ACB | 0.571 | 4.20 | 3.32 | 20.84 | 0.601 |
|  | HG02343 | ACB | 0.585 | 4.19 | 3.35 | 20.17 | 0.615 |
|  | HG02420 | ACB | 0.595 | 4.12 | 3.29 | 20.22 | 0.626 |
|  | HG01988 | ACB | 0.603 | 4.04 | 3.26 | 19.37 | 0.632 |
| **mildly affected** | NA19189 | YRI | 0.787 | 4.26 | 3.98 | 6.67 | 0.871 |
|  | HG00232 | GBR | 0.814 | 3.48 | 3.26 | 6.13 | 0.905 |
|  | HG00530 | CHS | 0.824 | 3.49 | 3.30 | 5.49 | 0.919 |
|  | HG00542 | CHS | 0.827 | 3.49 | 3.29 | 5.60 | 0.923 |
|  | HG00531 | CHS | 0.839 | 3.49 | 3.32 | 4.86 | 0.938 |
|  | NA18960 | JPT | 0.840 | 3.55 | 3.36 | 5.22 | 0.947 |
|  | NA18856 | YRI | 0.862 | 4.31 | 4.16 | 3.49 | 0.965 |
|  | HG00116 | GBR | 0.865 | 3.51 | 3.38 | 3.66 | 0.969 |
| **reference** | NA12878 | CEU | 0.885 | 3.52 | 3.44 | 2.24 | 0.989 |
|  | average | ACB (n=89) | 0.887 | 4.26 | 4.17 | 2.02 | 0.988 |
|  | st.dev. | ACB (n=89) | 0.0024 | 0.03 | 0.04 | 0.13 | 0.002 |
|  | average | all (n=2487) | 0.885 | 3.74 | 3.66 | 2.19 | 0.988 |
|  | st.dev. | all (n=2487) | 0.0028 | 0.33 | 0.32 | 0.17 | 0.002 |

To evaluate the nature of these discrepancies, we tabulated the number of biallelic autosomal SNVs observed for each individual genome in each of the seven datasets. For most individuals we observed about 2% fewer SNVs in TGP38X than in TGP37 (Table 2 and Figure 3). Possible explanations for this reduction might include: changes in the reference (including reference/alternate allele switches), improved variant calling leading to fewer false positives, and stricter coverage requirements leading to more false negatives. The eight mildly affected genomes have 3.5%-6.7% fewer SNVs in TGP38X than TGP37. In contrast, the eight strongly affected genomes have 20-22% fewer SNVs in TGP38X than in TGP37. These high missingness values are well beyond the range observed for the mildly affected genomes (5.14% +/- 1.11%), which are themselves well beyond the range of the majority of genomes (2.19% +/- 0.17%). While the strongly affected ACB genomes have similar SNV counts in TGP37 to the remaining 89 ACB individuals, they have markedly fewer SNVs in TGP38X. We also observed reduced genotype concordance for the strongly affected genomes relative to the other ACB individuals (Table 2). Finally, comparison with the TGP38H and TGP38N versions, which were constructed using an independent set of sequence reads, shows that the SNV counts in these individuals are concordant between TGP37, TGP38H and TGP38N (Figure 3). Taken together, these differences suggest that TGP38X erroneously excludes a set of 0.8-0.9 million SNVs from the strongly affected ACB genomes; these SNVs are included by independent analyses (e.g., TGP37 and TGP38N). TGP38S is similarly affected.

### Evaluation of the associated samples of related individuals

We evaluated the degree of relatedness of the 150 ‘related individuals’ in TGP38Xr (Figure 2B), expecting all of them to show some degree of relatedness to at least one of the individuals in the main TGP cohort -- TGP38X. (Due to the similarity between TGP38Sr and TGP38Xr, we elected to only report results for the latter.) We computed fingerprints for all TGP38Xr individuals, then compared them to all TGP38X individuals and to each other (Table S2). Over two thirds of the TGP38Xr individuals can indeed be recognized as closely related to TGP38X individuals, with fingerprint correlations > 0.45. On the other hand, at least 28 of the TGP38Xr individuals seem not to be related to anyone else in TGP38X or to each other (fingerprint correlation < 0.4).

One of the ‘related individuals’ in TGP38Xr (HG03982, from the Sri Lankan Tamil population in the UK, coded ‘STU’) has fingerprint correlation of 0.868 to an individual in TGP38X (HG03858, also STU). This fingerprint correlation level would suggest these are the same individual, and yet in IGSR, HG03858 is annotated as female while HG03982 is annotated as male. Neither individual has any relatives annotated in either TGP38X or TGP38Xr, nor could we identify any relatives by fingerprint comparison. We considered various hypotheses, including whether these individuals could be sex-discordant monozygotic twins (as a result of sex change, through differential resolution of XXY karyotype, mosaicism, etc.), the result of mislabeling of twin samples, or mislabeled, redundant samples of the same individual. TGP37 data support HG03858 being genetically female, with two copies of chrX and no chrY. To test these hypotheses, we evaluated whether HG03982 could indeed be a male sample, as annotated.

1. No chrY variant calls were released for TGP38Xr, and chrX variant calls are available only in the pseudoautosomal regions (PARs, which combine data from chrX and chrY). We computed chromosome-specific fingerprints, including for the PARs. The resulting 0.928 correlation of PAR-specific fingerprints of HG03858 and HG03982 suggests these two samples have the same sex-chromosome karyotype (XX) and the same chrX haplotypes, consistent with being the same individual, or identical twins. For comparison, we observe 0.954 correlation of PAR-specific fingerprints of samples HG00578 and HG00635 (female siblings, with overall autosomal fingerprint correlation of 0.689), and 0.622 correlation of samples HG00512 and HG00501 (male and female siblings, respectively, with overall autosomal fingerprint correlation of 0.687).
2. We observed 91.8% genotype concordance in the PARs of HG03858 and HG03982, consistent with these being the same person or female siblings. For comparison, the PAR genotype concordance of female siblings HG00578 and HG00635 was 94.1%, and the PAR genotype concordance of male and female siblings HG00512 and HG00501 was 45.1%.
3. Based on the coverage levels along chrX and chrY (from low-coverage data), we estimated the number of chrX and chrY copies for all STU samples and found that HG03982 clusters with the female samples (Figure S1A).

We conclude that HG03858 and HG03982 are both genetically female. Lacking further information about the individuals, we hypothesize that these two samples derive from the same individual or from identical twins, and that HG03982 may have been annotated as male as a result of a clerical error.

When comparing, through fingerprinting, the partially overlapping sets of related individuals in TGP38Xr and TGP38Nr (Figure 2B), we identified pairs of individuals with different identifiers but with genome fingerprint correlations consistent with them being the same individual, suggesting identifier swaps: genome HG03797 in TGP38Xr is equivalent to genome HG03799 in TGP38Nr, and conversely genome HG03799 in TGP38Xr is equivalent to genome HG03797 in TGP38Nr. Since HG03797 and HG03799 are, respectively, the father and mother in different trios (Bengali families BD16 and BD17), we were able to determine, through fingerprint comparisons with the offspring in these trios, that the genome identifiers were inadvertently swapped in TGP38Xr (and similarly in TGP38Sr). We similarly observed a potential identifier swap involving genomes HG03699 and HG03700, which are respectively the father and mother in Punjabi family trio PK59. In this case, fingerprint comparisons to the offspring could not elucidate the nature of the discrepancy, since both parents are equally related to their child. In this case, we used estimated chrX/chrY copy numbers in the TGP38Nr data, as described above, to determine that HG03699 is male and HG03700 is female, consistent with the metadata on these individuals (Figure S1B). Thus, we conclude that the identifiers were again mistakenly swapped in TGP38Xr (and TGP38Sr).

### Evaluation of fingerprint differences across TGP versions

While comparison of results pertaining to a high-quality genome (NA12878) can validate that a data processing pipeline has the capacity to perform its intended task well, it cannot assess whether the pipeline performs consistently across different input data. Since in-depth comparison of every genome would be low-throughput and expensive, we used genome fingerprints for their primary purpose: to perform rapid “self-comparison” of every genome across the different processing pipelines (Figure 4).

The comparisons among TGP37, TGP38L and TGP38C (Figure 4, columns 1-3) corresponded well for every genome (⍴ > 0.96). In addition, the low variability in fingerprint correlation, both within and across TGP populations (colors), suggests that the liftOver and CrossMap pipelines perform very consistently across individuals.

The TGP38S and TGP38X versions of each genome differ by whether indels were reported as variants (TGP38X only) along with the SNVs. For the majority of genomes, self-comparisons between these versions (Figure 4, column 12) corresponded well and varied little across genomes (⍴ > 0.97). However, a subset of 17 genomes (‘strongly affected’ in Table 3) show very reduced fingerprint correlations, very reduced SNV counts, and low genotype concordances. These include 9 genomes from the ACB population, adding HG02436 (which is absent from the TGP37 set) to the 8 ACB genomes described above. A further set of 15 individuals (‘mildly affected’ in Table 3) showed an intermediate, outlying level of degradation in self-correlations, SNV counts, and genotype concordance. Since these same 32 individuals include the outliers in all other comparisons involving versions TGP38S or TGP38X (Figure 4, columns 7-17), while the comparisons not involving these TGP versions do not produce comparable outliers (Figure 4, columns 1-6 and 18-21), we deduce that the pipelines used to create the TGP38S and TGP38X versions were less successful in processing these genomes than the rest of the cohort.

**Table 3.**
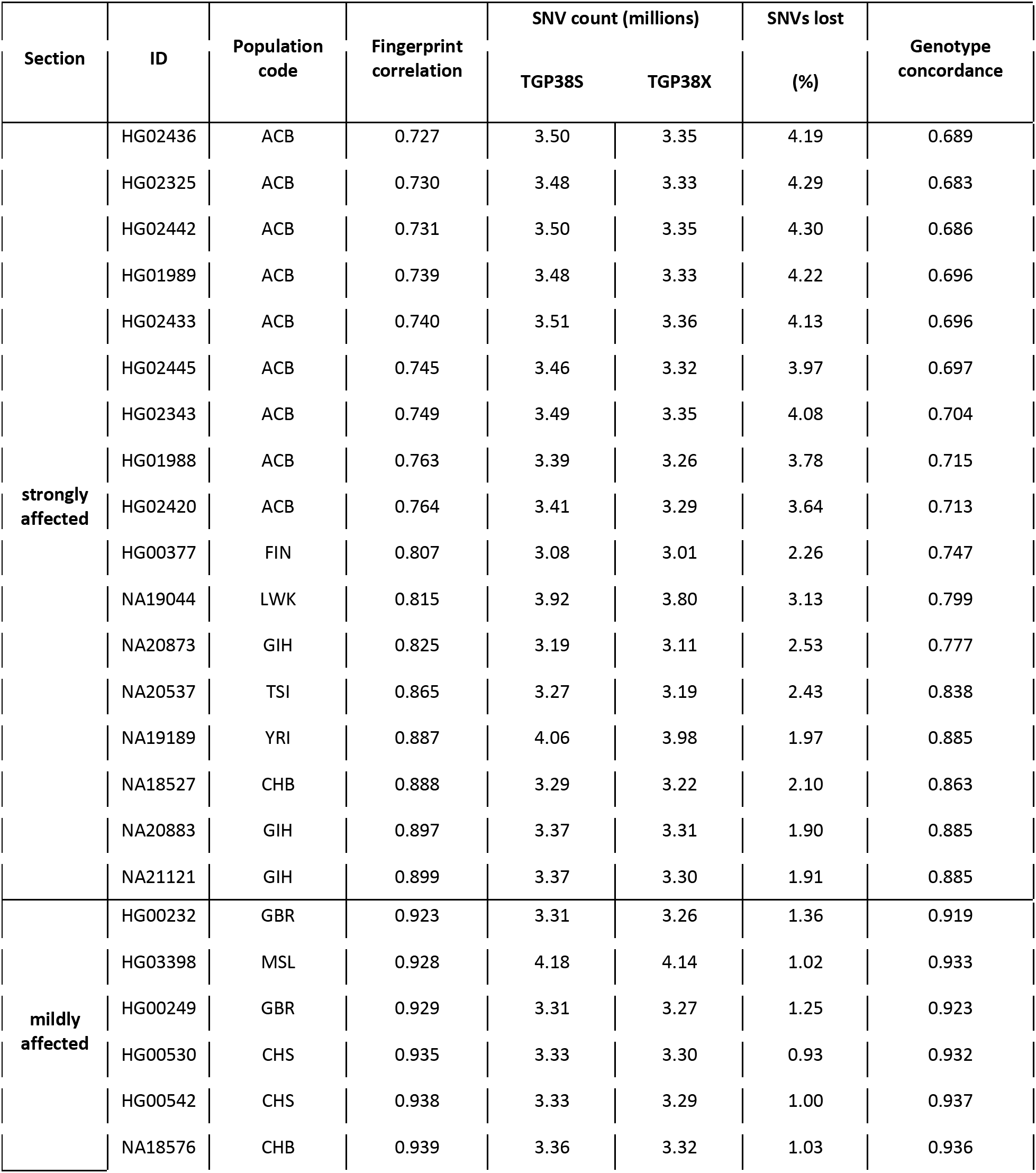
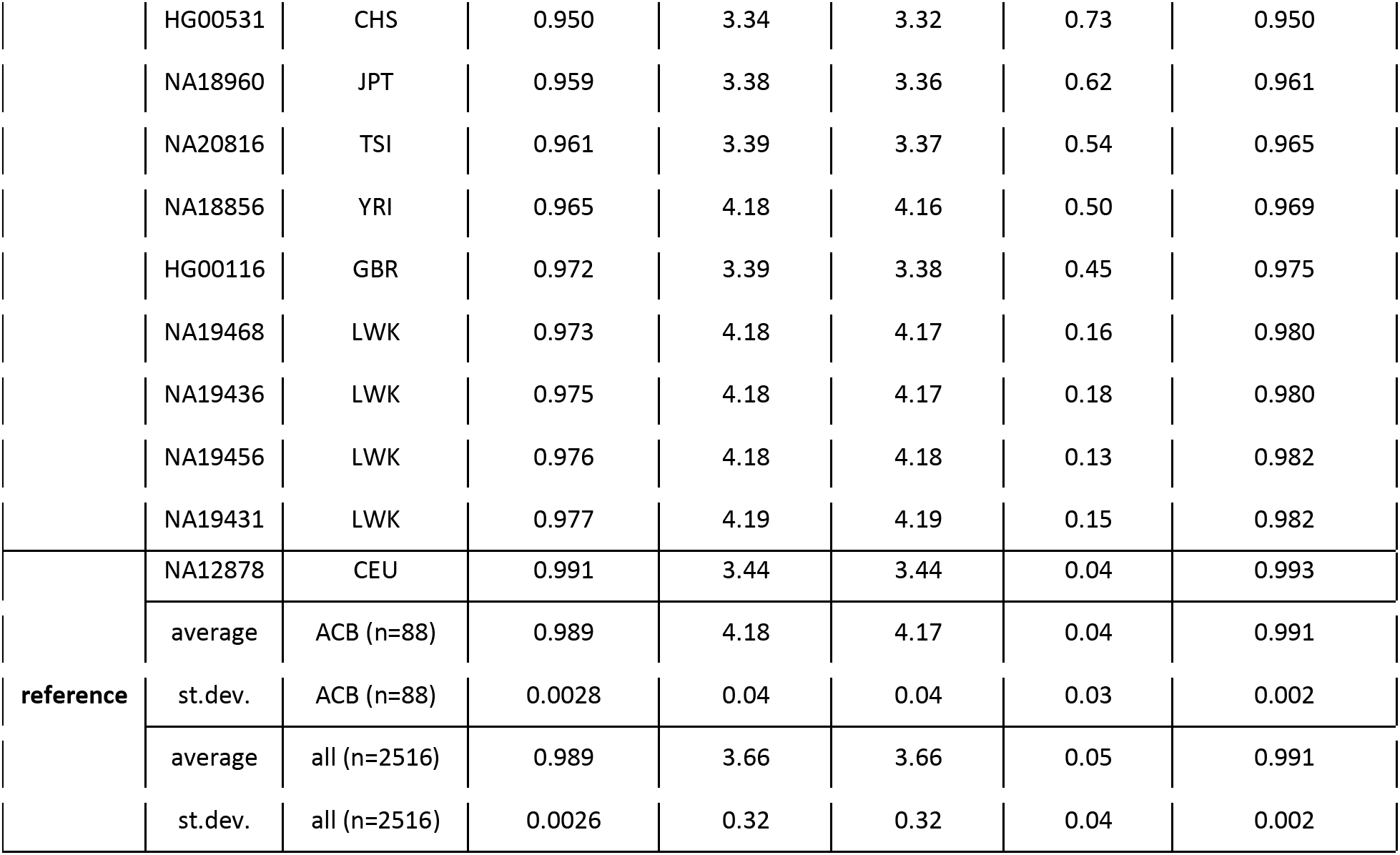
Observed statistics for the outlier individuals most affected by variant calling including or excluding indels (TGP38S vs. TGP38X), in comparison to the ‘platinum’ NA12878 genome, the 88 ACB individuals unaffected by this bioinformatic difference, and the 2516 similarly unaffected individuals in the entire cohort.

The TGP38H versions of each genome differ by inclusion of many additional variants, particularly around centromeric regions (see next section). This leads to reduced self-correlations in comparisons between TGP38H and other TGP versions (Figure 4, columns 16- 21).

### Identification of genomic regions with excess variation, or missing variation

The various TGP versions differ significantly in the total number of SNVs reported. We compared in pairwise fashion the four versions of the TGP expressed relative to GRCh38, by tallying (from the VCFs) how many variants are shared (i.e., both datasets have a SNV at a given coordinate, with the same REF and ALT alleles), and how many are unique to each set. We based this analysis on the 2,503 genomes shared by all TGP versions (i.e., all genomes in the original TGP37 set, minus NA18498). We found that each TGP version has a large number of SNVs absent from the other versions (Table 4), and that the chromosomal distribution of these discrepancies is a combination of (1) a nearly uniform background level characteristic to each dataset and (2) multiple clusters of variants unique to one dataset (Figure 5).

**Table 4.** Statistics of pairwise comparisons of the four GRCh38 callsets. Minimum, maximum, average and standard deviation of all fingerprint self-correlations, number of shared SNVs, number of SNVs unique to each set, and average genotype concordance over the 2,503 individuals included.

| Set 1 | Set 2 | Fingerprint self-correlation |  |  |  | Shared SNVs | Unique SNVs |  | Genotype concordance |
| --- | --- | --- | --- | --- | --- | --- | --- | --- | --- |
|  |  | min | max | avg | std |  | to Set 1 | to Set 2 |  |
| TGP38L | TGP38S | 0.601 | 0.897 | 0.889 | 0.015 | 68,021,487 | 9,344,300 | 3,407,548 | 0.959 |
| TGP38L | TGP38X | 0.552 | 0.897 | 0.888 | 0.018 | 68,079,268 | 9,291,699 | 3,720,873 | 0.959 |
| TGP38S | TGP38X | 0.727 | 0.996 | 0.988 | 0.017 | 71,334,231 | 123,182 | 185,835 | 0.971 |
| TGP38L | TGP38H | 0.792 | 0.829 | 0.812 | 0.006 | 74,767,408 | 2,605,724 | 29,944,460 | 0.944 |
| TGP38S | TGP38H | 0.542 | 0.817 | 0.800 | 0.015 | 68,970,797 | 2,473,081 | 35,748,569 | 0.954 |
| TGP38X | TGP38H | 0.500 | 0.817 | 0.799 | 0.017 | 69,028,474 | 2,478,010 | 35,690,845 | 0.951 |

A particularly noteworthy signal involves a very large number of SNVs in and around centromeric regions, unique to TGP38H (Figure 5 lower inset, and Figure S2). Their absence in TGP38L and TGP38C is not surprising, since GRCh37 lacked centromeric sequence models and thus there were no centromeric variants in TGP37 to be lifted over; as for TGP38S and TGP38X, the method used for generating these versions filtered out variants in centromeres.^6^ The density of centromeric variants in TGP38H also significantly exceeds the density observed elsewhere along the chromosomes (Figure S2).

Through pairwise comparison of the TGP versions, we identified 104,572 genomic segments significantly enriched (or depleted) in SNVs unique to one TGP version relative to the other (see examples in Figure 5 insets) and evaluated them for overlap with genomic features. We found that the segments with the largest counts of unique variants correspond to centromeric, telomeric and other heterochromatin and satellite sequences. After excluding these segments, we merged the 95,843 remaining ones by overlap into 12,307 regions.

The longest of these regions was a gene-dense 2.96 Mb segment on chromosome 12 erroneously lacking variation in TGP38L (Figure 5, middle inset). This segment (chr12:6889106- 9847458, between rs143703503 and rs111979444) has 71,946 variants unique to TGP38X (112,750 in TGP38H) but no variants shared with TGP38L, and no SNVs unique to TGP38L. In fact, the only variation reported in this region in TGP38L includes two indels (rs11394173 and rs56027063) with alternate allele frequency of 1, suggesting an error in the reference sequence. On the other hand, the corresponding range in GRCh37 (chr12:6999223-10000058) includes 43,406 variants (SNVs and indels) in TGP37. We verified that it is possible, using the liftOver tool, to transform coordinates in this region from GRCh37 to GRCh38. We conclude that a bioinformatic pipeline error likely caused this extensive set of variants to fail to lift over from TGP37 to TGP38L, yielding a significant gap in the reference variation. We note that this is a large and gene-dense region, including 100-150 transcript clusters, 75 of which are not pseudogenic. Bioinformatic analyses in this region, using annotated variation in TGP38L as reference, may have been confounded by this issue.

We evaluated the remaining regions over 250 kb in length and found them to largely correspond to genomic loci that were missing (gaps) or misassembled in previous versions of the reference genome (Table S3). Most of these regions have since been corrected or partially improved in GRCh38, and some still include gaps. Of note, since much knowledge on population variation was annotated (and submitted to dbSNP) based on older genome reference versions, and then lifted over to the current version, many of these regions show significantly reduced variation compared to the regions surrounding them, e.g., when visualized via genome browsers. For example, when comparing TGP38L to TGP38X we identified a 135 kb segment on chromosome 6 with 4,588 SNVs unique to TGP38X, but no variation in TGP38L. These 135 kb of sequence in GRCh38 replace a 50-kb clone gap in GRCh37 (GRC issue HG-564). As a result, this region is significantly depleted in variants in dbSNP (Figure S3).

### Identification of alternate allele discrepancies

One of several improvements in GRCh38 relative to GRCh37 was the correction of several thousand reference alleles from a minor/rare allele to the major allele ^20^. Since TGP38L, TGP38C, TGP38S, TGP38X, TGP38H and TGP38N are all represented relative to the same reference sequence, all variants shared between any two of these TGP versions have the same reference allele. On the other hand, while tallying shared and unique variants among the TGP versions (previous section), we identified SNVs with discrepant alternate (ALT) alleles. While TGP38S and TGP38X had the same ALT allele throughout the genome, and there were few ALT differences between TGP38L and TGP38C (both being directly derived from TGP37), we observed (tens of) thousands of discrepant ALT alleles genome-wide between every other pair of TGP versions (Table 5). These discrepancies are enriched in exons and in coding regions, relative to the number expected from the densities observed in non-exonic and non-coding regions, respectively, perhaps due to the inclusion of exome-derived data in all TGP versions except TGP38H and TGP38N. The enrichment in coding regions is most prominent in comparisons involving TGP38S or TGP38X.

**Table 5.** Discrepant alternate (ALT) alleles of pairwise comparisons of the six GRCh38 callsets. All: number of autosomal variants (SNVs and indels) with different stated ALT alleles. AC>0: number of ALT-discrepant autosomal SNVs with allele counts of at least 1 in both datasets. AC>9: number of ALT-discrepant autosomal SNVs, further requiring the sum of the allele counts in both datasets to be at least 10. Exonic: number of AC>0 ALT-discrepant autosomal SNVs mapped to any exons (coding and non-coding). Coding: number of AC>0 ALT-discrepant autosomal SNVs mapped to coding exons. Observed/expected (exonic, coding): ratio of AC>0 ALT-discrepant SNVs in exons and coding regions, respectively, relative to their corresponding expected values from non-exonic and non-coding densities.

| Comparison | ALT change |  |  |  |  | observed/expected |  |
| --- | --- | --- | --- | --- | --- | --- | --- |
|  | All | AC>0 | AC>9 | exonic | coding | exonic | coding |
| L - C | 184 | 184 | 103 | 16 | 6 | 1.70 | 2.31 |
| L - N | 75729 | 74471 | 8122 | 3828 | 1174 | 0.97 | 1.10 |
| C - N | 75942 | 74696 | 8154 | 3852 | 1169 | 0.97 | 1.09 |
| S - N | 74137 | 71845 | 7126 | 4268 | 1571 | 1.13 | 1.53 |
| X - N | 73690 | 71691 | 6952 | 4260 | 1565 | 1.13 | 1.53 |
| L - S | 28351 | 28323 | 3484 | 1614 | 549 | 1.08 | 1.35 |
| C - S | 28397 | 28227 | 3418 | 1605 | 539 | 1.08 | 1.33 |
| S - X | 0 | 0 | 0 | 0 | 0 | NA | NA |
| L - X | 28358 | 27886 | 3105 | 1598 | 547 | 1.09 | 1.37 |
| C - X | 28328 | 27807 | 3047 | 1590 | 537 | 1.08 | 1.35 |
| S - H | 13535 | 9111 | 3899 | 459 | 190 | 0.95 | 1.46 |
| X - H | 13852 | 9012 | 3871 | 451 | 190 | 0.94 | 1.47 |
| L - H | 21106 | 17693 | 4989 | 876 | 307 | 0.93 | 1.21 |
| C - H | 21484 | 18114 | 5059 | 932 | 312 | 0.97 | 1.20 |
| H - N | 7818 | 6998 | 2004 | 220 | 102 | 0.58 | 1.01 |

Several reasons can contribute to the identification of discrepant alternate alleles. From observational noise alone, one can expect different alternate alleles to be observed at low frequencies, particularly for independently sequenced samples. Some ALT discrepancies can be partially explained by the fact that TGP38S and TGP38X report only biallelic sites, while TGP38L, TGP38C, TGP38H and TGP38N also include multiallelic sites. For example, at chr1:44,819,962 (chr1:45,285,634 in GRCh37 coordinates), TGP38L reports 3 individuals with ALT = A, while both TGP38S and TGP38X report 1 individual with ALT = C. In contrast, TGP38H reports a triallelic site with 3 individuals with ALT = A, and 1 with ALT = C. In some other cases, the discrepancy can be attributed to different representations of variation in the vicinity of insertions, deletions and multi-nucleotide variants. We thus tallied all ALT discrepancies, classified them based on these features (see Methods), and visualized the three resulting classes relative to their allele counts (Figure 6).

While most of the ALT-discrepant variants are rare (a fraction have AC = 0 in one or more of the TGP versions, see Table 5), some are high frequency and thus cannot be explained by observational noise. In fact, for all classes of discrepant ALT allele sites we observed loci where every individual is reported to be homozygous for a non-reference allele, but for different alleles in the two TGP versions.

Multiallelic sites in TGP38C discrepant with TGP38N span the frequency spectrum; discrepant sites that are biallelic in both versions but multiallelic in some other version (TGP38L, TGP38S, TGP38X or TGP38H) display generally higher frequencies in TGP38C than in TGP38N (Figure 6A). Indel-related variants are enriched in high frequencies and tend to have higher reported frequency in TGP38C relative to TGP38N (Figure 6B). Similarly, we observed a tendency for higher reported frequencies in TGP38C for simple biallelic SNVs, unaffected by indels, that are ALT discrepant with TGP38N (Figure 6C).

When comparing TGP38X to TGP38N, we observed a strong tendency for higher allele frequencies in TGP38N relative to TGP38X for multiallelic SNVs (Figure 6D) and those affected by indels (Figure 6E). We observed discrepant biallelic SNVs unaffected by indels at all frequencies (Figure 6F). We also noted an enrichment in discrepant biallelic SNVs with high frequencies in TGP38N but lower frequencies in TGP38X, potentially as a result of undercalling in the latter version.

### Analysis of phasing concordance

We evaluated the concordance among the six TGP versions that are phased (TGP37, TGP38L, TGP38C, TGP38S, TGP38X and TGP38N) in two ways: through phasing-aware fingerprinting, and through a phasing accuracy metric we described previously.^21^

First, we modified the procedure for computing genome fingerprints to capture phasing information. Genome fingerprints normally capture the presence of alternate alleles but ignore whether the alternate allele was phased relative to other variants. We modified the method to consider whether the alternate alleles in two consecutive heterozygous variants are phased to be on the same haplotype (e.g., both maternal) or on opposite haplotypes (i.e., one maternal and one paternal). We then encode the latter scenario (opposite haplotypes) by swapping the reference and the alternate alleles for the second variant. In this way, phasing-aware genome fingerprints can capture local phasing information (microhaplotypes), leading to decreased similarity between genomes that differ in such local phasing.

We computed phasing-aware genome fingerprints for the six phased TGP versions and evaluated the change in self-correlation for each individual. When comparing versions TGP37 and its liftover derivatives TGP38L and TGP38C (Figure 7A, columns 1-3), we observed that adding phasing information leads to slightly higher self-correlations compared to standard fingerprints (Figure 4), as expected from consistent phasing. In contrast, in all other comparisons adding phasing information strongly reduces correlations, indicating phasing discrepancies between the TGP versions. This reduction in correlations is generally stronger for genomes of African individuals. In comparisons involving either TGP38S or TGP38X to other versions, we again observed several outliers with reduced correlations (including the degraded ACB genomes). In contrast, five genomes have much higher correlations than the rest, suggesting their phasing was much less inconsistent across versions: NA12878, NA10851, NA10847 and NA07048 from the CEU population, and NA19129 (YRI). An additional four genomes also have increased correlations when comparing TGP38S or TGP38X to TGP37, TGP38L or TGP38C: NA20910 (GIH), NA19445 (LWK), HG00867 (CDX), and HG00155 (GBR).

When comparing TGP38S and TGP38X (Figure 7A, column 12), we observed overall degradation indicating phasing discrepancies between these two related TGP versions: there were many outliers (including the degraded ACB genomes), and one third of the genomes had much stronger discrepancies in phasing. Surprisingly, this analysis revealed a differential effect by population: the most affected genomes include all the individuals in the SAS populations BEB, ITU, PJL and STU, but none from GIH; all the individuals in the AFR populations ESN, GWD and MSL, but none from LWK and YRI; sizable subsets of AMR populations ACB, CLM, PEL and PUR; and a few individuals from EAS populations CDX and CHS, EUR populations FIN, GBR and IBS, and AMR populations ASW and MXL.

Second, we studied the VCFs of phased TGP versions on GRCh38 in pairwise fashion to identify phase switches (one version relative to the other) in each individual. Using a phasing accuracy metric,^21^ we computed for each individual the level of phasing agreement between TGP38L and TGP38X, and separately between TGP38S and TGP38X (Figure 7B). We observe overall very low phasing concordance between TGP38L and TGP38X except for the same nine individuals highlighted in the fingerprint-based results. Similarly, the TGP38S-TGP38X comparison shows differential phasing consistency per population (see population annotations in Figure 7B), again corroborating the fingerprint-based results.

## Discussion

We presented here the application of genome fingerprints^11^ for quick and simple comparison of seven versions of the Thousand Genomes Project (TGP) dataset, expressed relative to two versions of the reference genome (GRCh37 and GRCh38); the ability to compare genomes across different versions of the reference sequence was a specific goal in the design of genome fingerprints. In addition to the overall comparison of the full datasets of phased SNVs and indels (TGP37 vs. TGP38X, and related samples), other pairwise comparisons of TGP versions provided insights into the effects of lifting over variants from one reference version to the other (TGP37 vs. TGP38L/C), of lifting over vs. native mapping and variant calling (TGP38L/C vs. TGP38X), of different variant calling procedures (TGP38X vs. TGP38S), and of high-coverage sequencing (TGP38H/N). Through these comparisons, we identified multiple discrepancies between the datasets, pointing at changes in the list of included genomes, some additional cryptic relationships, overall changes in biallelic SNV counts, more significant changes in SNV counts and reduced genotype concordance affecting a subset of the individuals, and low concordance of variant phasing. These observations illustrate the value of employing data reduction techniques, such as the genome fingerprints used here, to enable quality evaluation of the output from every input to an automated pipeline, and not just a “gold-standard” subset.

Minor reference alleles are a problem for identifying and reporting clinical variants^22, 23^, and discrepant ALT alleles could produce similar issues. Cross-comparison of TGP versions identifies many discrepancies of alternate alleles observed. While some reflect infrequent observations (low allele counts), many discrepantly observed alleles seem to be very frequent in the population. There can be multiple causes for ALT allele discrepancies, including the presence of multiple alleles, and overlap or vicinity with indels and more complex variants. In many cases, these can only be identified through comparison with an additional dataset. For example, an ALT-discrepant biallelic SNV between TGP38C and TGP38X may overlap by coordinate with an indel reported in TGP38N. This suggests that there is still room for improvement in variant-calling and variant-reporting pipelines, particularly in the presence of multiple alleles, indels, and low complexity sequences. Differences in reference sequences were shown to lead to discrepancies in variant calls in the exome^24^. We observed these discrepancies to be genome-wide, with an enrichment in ALT allele discrepancies in coding regions, particularly when comparing TGP versions based on a mixture of exome and low-pass whole-genome sequencing vs. versions based solely on 30x whole-genome sequencing. This suggests there may still be some difficulty integrating the two data types, and that less biased results may be obtained from comprehensive whole-genome sequencing, particularly using long reads^25^.

We evaluated the consistency of phasing among TGP versions using two methods: phasing-aware fingerprinting and a phasing accuracy metric, which respectively emphasize high-frequency or low-frequency switch errors.^21^ These comparisons revealed significant discrepancies in phasing among the different TGP versions for most (but not all) genomes. While pairwise comparisons can reveal discrepancies, it is not always evident which one is the correct result. In the context of phasing, it is reasonable to assume that TGP38N is the better phased version, since its phasing method included multiple related individuals.

Notably, a small subset of genomes showed more consistent phasing; these genomes include those most commonly studied and used for evaluating methodologies. Current best practices for benchmarking variant calls are largely based on the use of ‘truth set’ resources of the Genome In A Bottle (GIAB) Consortium.^12–14^ Specifically, TGP38X was evaluated by comparing the variant call sets observed for the ‘platinum’ NA12878 genome, and computing false positive and false negative call rates in regions for which the GIAB considers calls to be high confidence.^6^ Strathern’s generalization of Goodhart’s Law, “When a measure becomes a target, it ceases to be a good measure,”^26^ would seem to apply; since the Platinum NA12878 genome and other GIAB ‘truth sets’ are used to train experimental and computational technologies, it seems inappropriate to also use them for QC purposes. We observe that such verification may be insufficient for global evaluation of large genome datasets including samples from diverse population backgrounds, which may be differentially affected by reference and software changes.

As a partial way to mitigate this deficiency, we recommend performing global dataset comparisons using genome fingerprints, and other general-purpose^27^ or domain-specific metrics. Such ‘relative benchmarking’, in which each individual genome can serve as its own reference, can supplement ‘absolute benchmarking’ relative to truth sets. As a result of such relative benchmarking, multiple discrepancies may become evident that cannot be immediately resolved in the absence of a truth set; resolving such discrepancies would certainly necessitate further computational analyses and, in some cases, experimental testing.

For most researchers, though, the existence of multiple versions of the TGP dataset raises an important question: ‘what version should I use?’ The answer will naturally depend on the specific analysis goals and available data. For studying genomes expressed relative to GRCh37, TGP37 is the obvious choice. For genomes expressed relative to GRCh38, though, there are multiple options available, differing from each other in multiple ways (Table 1). Since the beginning of this comparison work, version TGP38L was formally withdrawn, leaving TGP38C as the version most similar to TGP37 in content, and thus perhaps yielding the most comparable results. Versions TGP38S and TGP38X are limited to biallelic content; we also found multiple quality issues in these versions. TGP38H is the only version that includes variants in centromeric regions, which may be an important benefit, or a negative, depending on the study. It is also by far the most difficult version to work with due to its size. Finally, TGP38N seems to have the most advantages, being based on comprehensive WGS resequencing of the original samples and phasing using families. We noted, though, that both TGP38C and TGP38N require some preprocessing due to their unusual representation of multiallelic variants and, in the case of TGP38N, the inclusion of 698 additional genomes that may not be needed for many studies.

## Description of Supplemental Data

Supplemental Data include three figures and three tables.

## Acknowledgments

This work was supported by NIBIB grant U54 EB020406 and NIA grant U19 AG023122.

## Web Resources

Project website: db.systemsbiology.net/gestalt/tgpqc

Genome fingerprints project website: db.systemsbiology.net/gestalt/genome_fingerprints

Genome fingerprints Github page: github.com/gglusman/genome-fingerprints

IGSR: International Genome Sample Resource, http://www.internationalgenome.org

liftOver tool: http://genome.ucsc.edu/cgi-bin/hgLiftOver

**TGP37**: http://ftp.1000genomes.ebi.ac.uk/vol1/ftp/release/20130502/

**TGP38L**: http://ftp.1000genomes.ebi.ac.uk/vol1/ftp/release/20130502/supporting/GRCh38_positions/

**TGP38C**:

http://ftp.1000genomes.ebi.ac.uk/vol1/ftp/data_collections/1000G_2504_high_coverage/working/phase3_l iftover_nygc_dir/

**TGP38S**:

http://ftp.1000genomes.ebi.ac.uk/vol1/ftp/data_collections/1000_genomes_project/release/20181203_bial lelic_SNV/

**TGP38X**:

http://ftp.1000genomes.ebi.ac.uk/vol1/ftp/data_collections/1000_genomes_project/release/20190312_bial lelic_SNV_and_INDEL/

**TGP38H**:

ftp://ftp.1000genomes.ebi.ac.uk/vol1/ftp/data_collections/1000G_2504_high_coverage/working/20190425

_NYGC_GATK/

**TGP38N** and **TGP38Nr**:

http://ftp.1000genomes.ebi.ac.uk/vol1/ftp/data_collections/1000G_2504_high_coverage/working/2020102 8_3202_phased/

**TGP37r**: http://ftp.1000genomes.ebi.ac.uk/vol1/ftp/release/20130502/supporting/related_samples_vcf/

**TGP38Sr**:

http://ftp.1000genomes.ebi.ac.uk/vol1/ftp/data_collections/1000_genomes_project/release/20181203_bial lelic_SNV/supporting/related_samples/

**TGP38Xr**:

http://ftp.1000genomes.ebi.ac.uk/vol1/ftp/data_collections/1000_genomes_project/release/20190312_bial lelic_SNV_and_INDEL/supporting/related_samples/

## Data and Code Availability

Genome fingerprints for all datasets are available through the genome fingerprints project website. Code for computing genome fingerprints is available from Github.

---

**Figure S1.**
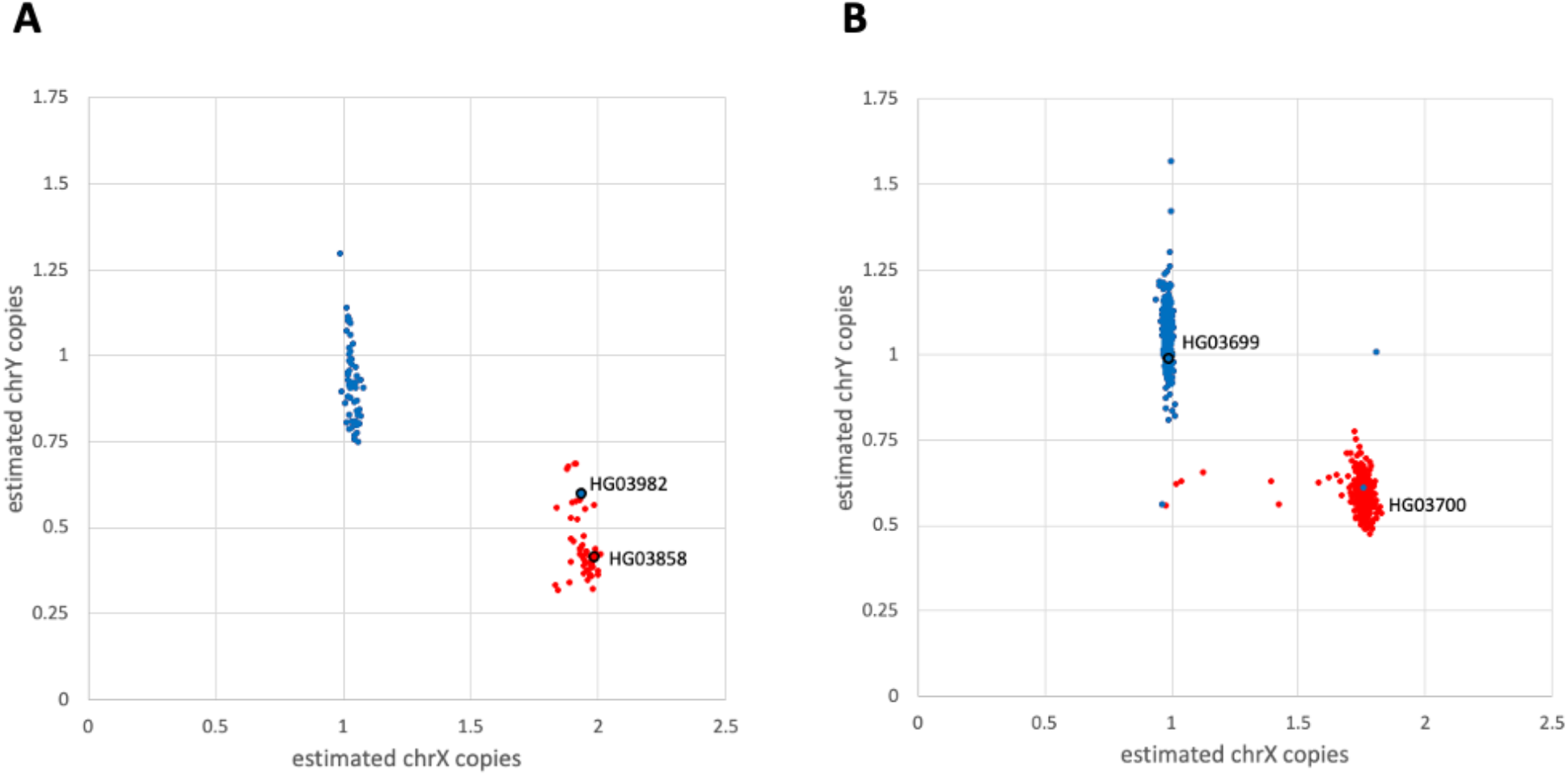
Estimated copy numbers for chromosomes X and Y. Red and blue points represent individuals annotated as female and male, respectively. A. Individuals in the STU population. HG03982, annotated as male, clusters with female samples. B. The 698 individuals in TGP38Nr. HG03699 and HG03700 cluster with male and female samples, respectively, consistent with annotation. (Other samples are seen with discrepant copy numbers.)

**Figure S2.**
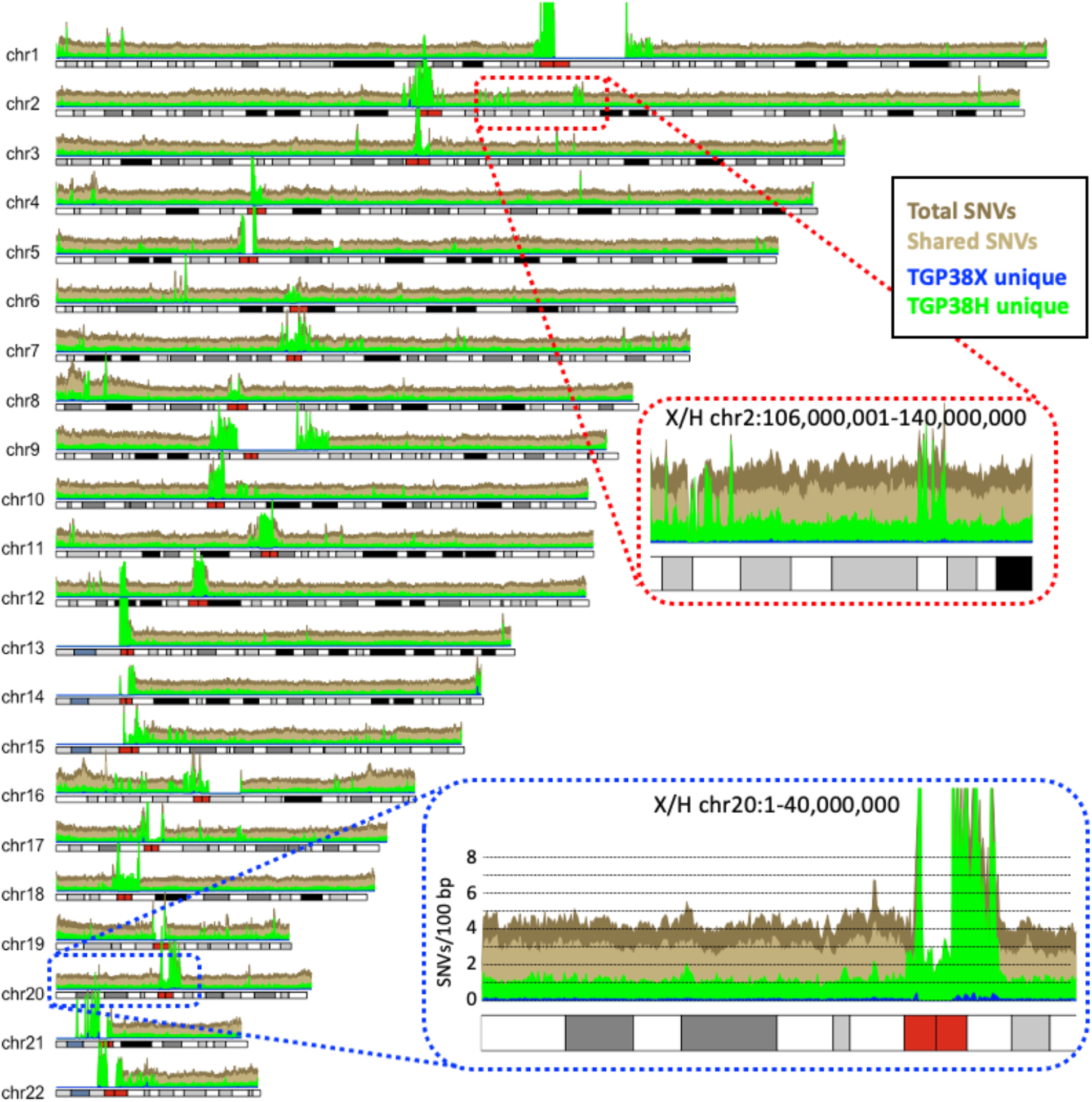
Density map of shared and unique variants along the chromosomes. The main plot shows the comparison between TGP38X and TGP38H, with shared variants in light brown, variants unique to TGP38X in blue, those unique to TGP38H in green, and the union of all variants in dark brown. Top inset (red outline): detail of a 36 Mb region on chr2 showing patches enriched in variants unique to TGP38H (absent from TGP38X). Lower inset (blue outline): detail of the first 40 Mb of chr20 highlighting the overall higher density of TGP38H-unique variants relative to TGP38X-unique variants, and the excess variation unique to TGP38H in centromeric regions.

**Figure S3.**
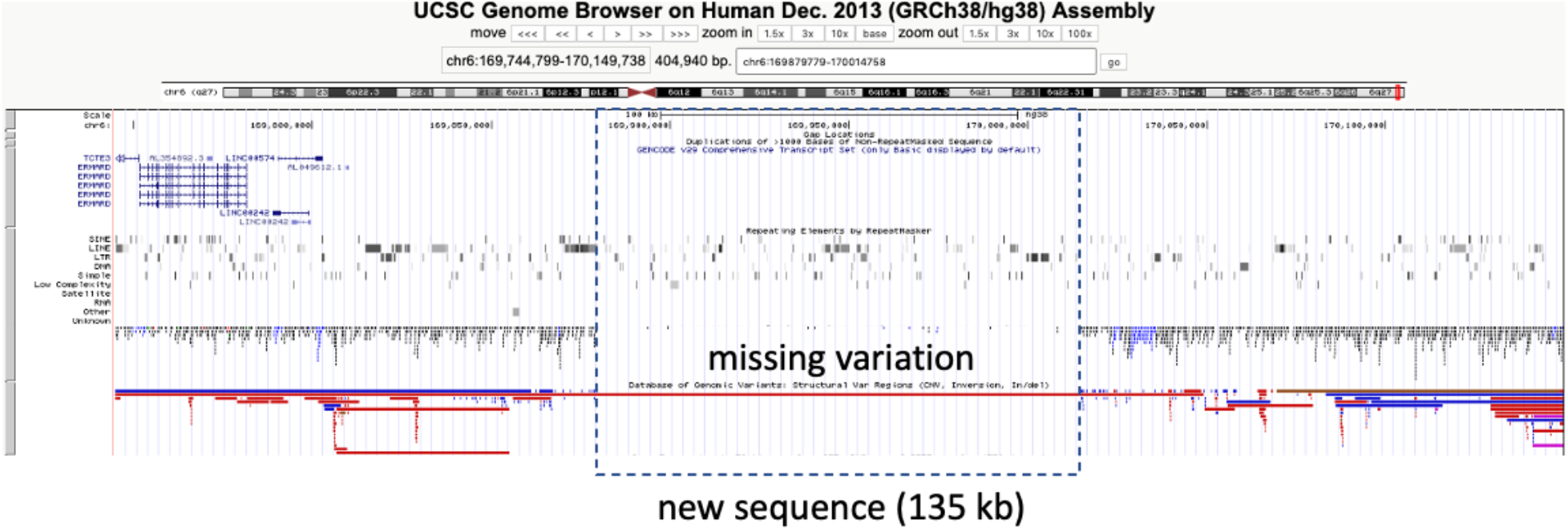
Example of a region missing variation in dbSNP. This segment on chromosome 6 includes 135 kb of new sequence that closed a former 50-kb clone gap. Many variants had been identified around the clone gap and submitted to dbSNP, but very few variants have been annotated in the new sequence since its release.

**Table S1.** The 45 additional individuals in TGP38X, absent from TGP37. None of these individuals have any annotated relationships in IGSR, but 34 have fingerprint correlations of at least 0.45 to other individuals in TGP38X, suggesting relatedness.

| ID | Population code | Most similar among individuals present in TGP37 |  | Most similar among TGP38X additional individuals |  |
| --- | --- | --- | --- | --- | --- |
|  |  | ID | Correlation | ID | Correlation |
| NA18576 | CHB | HG00530 | 0.521 | NA18527 | 0.515 |
| NA18527 | CHB | HG00530 | 0.514 | NA18576 | 0.515 |
| HG02358 | CDX | HG00530 | 0.506 | HG02405 | 0.498 |
| HG02170 | CDX | HG02152 | 0.506 | HG02173 | 0.498 |
| HG02168 | CDX | HG02396 | 0.505 | NA18576 | 0.503 |
| HG02169 | CDX | HG01797 | 0.505 | HG02173 | 0.496 |
| HG02173 | CDX | HG02389 | 0.502 | HG02168 | 0.502 |
| HG02176 | CDX | HG01802 | 0.502 | HG02170 | 0.497 |
| HG02405 | CDX | HG00844 | 0.501 | HG02358 | 0.498 |
| HG00134 | GBR | HG00142 | 0.499 | HG00249 | 0.460 |
| NA18791 | CHB | NA18960 | 0.498 | NA18576 | 0.496 |
| NA18955 | JPT | NA19005 | 0.497 | NA18576 | 0.493 |
| HG00377 | FIN | HG00329 | 0.481 | NA20873 | 0.492 |
| NA20873 | GIH | HG00232 | 0.457 | HG00377 | 0.492 |
| NA20537 | TSI | HG00232 | 0.469 | HG00377 | 0.491 |
| NA20816 | TSI | NA20516 | 0.482 | HG00377 | 0.472 |
| HG00249 | GBR | HG00232 | 0.481 | HG00377 | 0.482 |
| NA21121 | GIH | HG04001 | 0.467 | NA20883 | 0.480 |
| NA20883 | GIH | NA20854 | 0.462 | NA21121 | 0.480 |
| HG00152 | GBR | NA12287 | 0.467 | HG00377 | 0.470 |
| HG00270 | FIN | HG00269 | 0.466 | HG00377 | 0.469 |
| HG00104 | GBR | HG00157 | 0.461 | HG00377 | 0.468 |
| HG00302 | FIN | HG00373 | 0.466 | HG00377 | 0.468 |
| HG00312 | FIN | HG00308 | 0.467 | NA20537 | 0.467 |
| HG00303 | FIN | HG00269 | 0.467 | HG00249 | 0.466 |
| HG04301 | GBR | HG00306 | 0.463 | HG00377 | 0.464 |
| HG04303 | GBR | NA11930 | 0.464 | NA20537 | 0.464 |
| HG01471 | CLM | HG02266 | 0.463 | HG00377 | 0.453 |
| HG00359 | FIN | HG00266 | 0.463 | HG00377 | 0.463 |
| NA20831 | TSI | NA11881 | 0.457 | HG00249 | 0.463 |
| NA20829 | TSI | HG00232 | 0.463 | HG00249 | 0.462 |
| HG00135 | GBR | HG01516 | 0.462 | HG00249 | 0.458 |
| HG00156 | GBR | HG00150 | 0.458 | NA20537 | 0.461 |
| HG04302 | GBR | HG00337 | 0.458 | NA20816 | 0.459 |
| HG02436 | ACB | HG02433 | 0.422 | HG00377 | 0.404 |
| NA19359 | LWK | NA19309 | 0.389 | NA19044 | 0.331 |
| NA19044 | LWK | HG01989 | 0.373 | HG02436 | 0.365 |
| HG03398 | MSL | NA19189 | 0.349 | NA19044 | 0.344 |
| HG03393 | MSL | HG03376 | 0.348 | HG03398 | 0.330 |
| HG03431 | MSL | NA19189 | 0.336 | HG03398 | 0.330 |
| HG03171 | ESN | NA18520 | 0.336 | NA19044 | 0.323 |
| HG03549 | MSL | HG02643 | 0.334 | HG03398 | 0.335 |
| NA19371 | LWK | NA19452 | 0.332 | NA19044 | 0.333 |
| NA19398 | LWK | HG02442 | 0.330 | NA19044 | 0.332 |
| HG03462 | MSL | HG03397 | 0.330 | HG03398 | 0.329 |

**Table S2.** The 150 related individuals in TGP38Xr, sorted by similarity to TGP38X or to other TGP38Xr individuals, starting with the HG03982-HG03858 pair discussed in the text.

| ID | Population code | Most similar among individuals in TGP38X |  | Most similar among individuals in TGP38Xr |  |
| --- | --- | --- | --- | --- | --- |
|  |  | ID | Correlation | ID | Correlation |
| HG03982 | STU | HG03858 | 0.868 | HG03761 | 0.430 |
| HG00512 | CHS | HG00524 | 0.710 | HG00501 | 0.687 |
| HG00578 | CHS | HG00581 | 0.707 | HG00635 | 0.689 |
| HG00501 | CHS | HG00524 | 0.701 | HG00512 | 0.687 |
| HG02046 | KHV | HG02067 | 0.701 | HG00866 | 0.490 |
| HG00577 | CHS | HG00584 | 0.697 | HG00418 | 0.501 |
| HG02381 | CDX | HG02373 | 0.694 | HG00427 | 0.489 |
| HG00635 | CHS | HG00581 | 0.676 | HG00578 | 0.689 |
| NA20526 | TSI | NA20792 | 0.678 | NA07346 | 0.453 |
| HG03948 | STU | HG03673 | 0.674 | HG04053 | 0.441 |
| HG00983 | CDX | HG00978 | 0.665 | HG00866 | 0.511 |
| HG02372 | CDX | HG02371 | 0.659 | HG00866 | 0.502 |
| HG02024 | KHV | HG02026 | 0.659 | HG00427 | 0.495 |
| HG00702 | CHS | HG00657 | 0.656 | HG00866 | 0.496 |
| HG00866 | CDX | HG00867 | 0.645 | HG00427 | 0.515 |
| NA20898 | GIH | NA20886 | 0.635 | HG03723 | 0.444 |
| NA20871 | GIH | NA20868 | 0.631 | HG03723 | 0.452 |
| NA12891 | CEU | NA12878 | 0.626 | NA12892 | 0.464 |
| HG00153 | GBR | HG00158 | 0.624 | NA11993 | 0.465 |
| HG00733 | PUR | HG00732 | 0.623 | NA11993 | 0.443 |
| NA19685 | MXL | NA19661 | 0.623 | NA19660 | 0.611 |
| NA19675 | MXL | NA19678 | 0.616 | NA11993 | 0.446 |
| HG03715 | ITU | HG03713 | 0.616 | HG03723 | 0.448 |
| NA12892 | CEU | NA12878 | 0.612 | NA11993 | 0.474 |
| NA19660 | MXL | NA19664 | 0.513 | NA19685 | 0.611 |
| NA20336 | ASW | NA20334 | 0.603 | NA11993 | 0.350 |
| NA19373 | LWK | NA19374 | 0.600 | NA19150 | 0.334 |
| NA20341 | ASW | NA20289 | 0.597 | HG01274 | 0.331 |
| NA19470 | LWK | NA19443 | 0.596 | NA19469 | 0.538 |
| NA19396 | LWK | NA19397 | 0.588 | NA19444 | 0.330 |
| HG02363 | CDX | HG02353 | 0.586 | HG00427 | 0.500 |
| HG02377 | CDX | HG02250 | 0.579 | HG00866 | 0.504 |
| HG02387 | CDX | HG02386 | 0.579 | HG00866 | 0.502 |
| HG00418 | CHS | HG00531 | 0.507 | HG00427 | 0.575 |
| HG00427 | CHS | HG00530 | 0.519 | HG00418 | 0.575 |
| NA20322 | ASW | NA20321 | 0.574 | NA20361 | 0.338 |
| HG02388 | CDX | HG02375 | 0.572 | HG00866 | 0.498 |
| NA19444 | LWK | NA19434 | 0.570 | NA19453 | 0.446 |
| HG00124 | GBR | HG00119 | 0.567 | NA11993 | 0.457 |
| NA19985 | ASW | NA19713 | 0.554 | HG01274 | 0.344 |
| NA19240 | YRI | NA19239 | 0.543 | NA19150 | 0.340 |
| NA19469 | LWK | NA19443 | 0.416 | NA19470 | 0.538 |
| NA19453 | LWK | NA19445 | 0.537 | NA19444 | 0.446 |
| NA19381 | LWK | NA19380 | 0.404 | NA19382 | 0.533 |
| NA19382 | LWK | NA19380 | 0.453 | NA19381 | 0.533 |
| NA19313 | LWK | NA19331 | 0.530 | NA19373 | 0.333 |
| HG02344 | PEL | HG01961 | 0.530 | HG01995 | 0.496 |
| NA20893 | GIH | NA20895 | 0.528 | HG03723 | 0.460 |
| HG01983 | PEL | HG01936 | 0.521 | HG02344 | 0.494 |
| HG01995 | PEL | HG01961 | 0.519 | HG02344 | 0.496 |
| NA20361 | ASW | NA20362 | 0.517 | HG01274 | 0.365 |
| HG02288 | PEL | HG01961 | 0.510 | HG01995 | 0.492 |
| HG00716 | CHS | HG00530 | 0.508 | HG00983 | 0.502 |
| HG02347 | PEL | HG01961 | 0.507 | HG02344 | 0.489 |
| HG02524 | KHV | HG00530 | 0.495 | HG00866 | 0.497 |
| NA11993 | CEU | HG00377 | 0.496 | NA12892 | 0.474 |
| HG02525 | KHV | HG00542 | 0.496 | HG00866 | 0.494 |
| HG01274 | CLM | HG00377 | 0.490 | HG01278 | 0.475 |
| HG04192 | BEB | HG03940 | 0.450 | HG04191 | 0.487 |
| HG04191 | BEB | NA21121 | 0.450 | HG04192 | 0.487 |
| HG01347 | CLM | HG01113 | 0.482 | HG01274 | 0.459 |
| HG03723 | ITU | HG00377 | 0.480 | NA20893 | 0.460 |
| HG01278 | CLM | HG00377 | 0.479 | HG01274 | 0.475 |
| NA19797 | MXL | HG02299 | 0.478 | HG02288 | 0.469 |
| NA07346 | CEU | HG00249 | 0.472 | NA12892 | 0.462 |
| HG01482 | CLM | NA19729 | 0.469 | HG01995 | 0.467 |
| HG04053 | ITU | HG04023 | 0.468 | HG04055 | 0.466 |
| HG04055 | ITU | HG04054 | 0.468 | HG04053 | 0.466 |
| HG00247 | GBR | HG00249 | 0.466 | NA07346 | 0.458 |
| HG04050 | ITU | HG03861 | 0.463 | HG04055 | 0.466 |
| HG03761 | PJL | NA21121 | 0.459 | HG03723 | 0.449 |
| HG03806 | BEB | NA20873 | 0.459 | HG03723 | 0.450 |
| HG04058 | ITU | NA21121 | 0.459 | HG03723 | 0.458 |
| HG04127 | STU | HG04038 | 0.458 | HG03723 | 0.450 |
| NA19737 | MXL | HG01961 | 0.458 | HG01278 | 0.448 |
| HG03606 | BEB | NA20883 | 0.457 | HG03723 | 0.447 |
| HG02218 | IBS | HG00377 | 0.456 | NA11993 | 0.457 |
| HG04174 | BEB | NA21121 | 0.456 | HG04058 | 0.447 |
| HG04114 | STU | NA20873 | 0.456 | HG03723 | 0.450 |
| HG04150 | BEB | NA20873 | 0.455 | HG00866 | 0.455 |
| HG03487 | PJL | NA21121 | 0.455 | HG03723 | 0.444 |
| NA19798 | MXL | HG00377 | 0.455 | NA12892 | 0.447 |
| HG03929 | BEB | HG04001 | 0.454 | HG03723 | 0.448 |
| HG04132 | BEB | NA20873 | 0.454 | HG04127 | 0.444 |
| HG03811 | BEB | NA21121 | 0.454 | HG03723 | 0.452 |
| HG01483 | CLM | HG00249 | 0.453 | NA12892 | 0.450 |
| HG01453 | CLM | HG02277 | 0.452 | HG01274 | 0.453 |
| HG03656 | PJL | NA21121 | 0.452 | HG03723 | 0.450 |
| HG03901 | BEB | NA21121 | 0.452 | HG03723 | 0.451 |
| HG04024 | ITU | NA20883 | 0.452 | HG03723 | 0.452 |
| HG03847 | STU | HG03689 | 0.452 | HG03723 | 0.451 |
| HG04147 | BEB | NA20883 | 0.452 | HG04150 | 0.441 |
| HG03972 | ITU | NA20883 | 0.452 | HG03723 | 0.448 |
| HG04128 | BEB | NA20873 | 0.452 | HG03723 | 0.451 |
| HG03842 | STU | NA21121 | 0.452 | HG03723 | 0.451 |
| HG02781 | PJL | HG03706 | 0.451 | HG03723 | 0.446 |
| HG04135 | BEB | NA21121 | 0.451 | HG03811 | 0.442 |
| HG03650 | PJL | NA20873 | 0.451 | HG04127 | 0.449 |
| HG03904 | BEB | NA21121 | 0.451 | HG04058 | 0.447 |
| HG04037 | STU | NA20873 | 0.451 | HG03723 | 0.450 |
| HG03988 | STU | NA21121 | 0.451 | HG03723 | 0.448 |
| HG03639 | PJL | HG02728 | 0.451 | HG03621 | 0.447 |
| HG01590 | PJL | NA20883 | 0.451 | NA20893 | 0.446 |
| HG03618 | PJL | HG00377 | 0.451 | HG03723 | 0.445 |
| HG01322 | PUR | HG00377 | 0.450 | NA11993 | 0.442 |
| HG04149 | BEB | HG03940 | 0.449 | HG03723 | 0.448 |
| HG03845 | STU | NA21121 | 0.449 | HG03723 | 0.448 |
| HG03621 | PJL | HG03019 | 0.448 | HG03639 | 0.447 |
| HG03794 | BEB | NA21121 | 0.447 | HG04058 | 0.442 |
| HG03700 | PJL | NA21121 | 0.444 | HG03723 | 0.440 |
| HG03633 | PJL | NA20883 | 0.444 | HG03723 | 0.439 |
| NA20313 | ASW | NA20282 | 0.443 | NA20361 | 0.334 |
| HG03699 | PJL | NA21121 | 0.441 | HG04150 | 0.432 |
| HG03799 | BEB | NA21121 | 0.440 | HG03797 | 0.436 |
| HG01195 | PUR | HG00377 | 0.439 | HG01274 | 0.438 |
| HG03797 | BEB | NA21121 | 0.438 | HG03799 | 0.436 |
| NA19311 | LWK | NA19027 | 0.438 | HG03309 | 0.330 |
| HG01480 | CLM | NA20537 | 0.435 | HG01278 | 0.438 |
| HG01477 | CLM | HG00377 | 0.433 | HG01278 | 0.436 |
| HG01473 | CLM | HG02277 | 0.435 | HG01274 | 0.434 |
| HG03383 | MSL | HG03391 | 0.432 | HG03508 | 0.323 |
| HG01452 | CLM | HG01961 | 0.430 | HG01983 | 0.428 |
| HG03454 | MSL | HG03457 | 0.389 | NA19150 | 0.335 |
| NA20344 | ASW | NA20340 | 0.371 | HG01274 | 0.340 |
| HG03574 | MSL | HG03556 | 0.370 | HG03373 | 0.330 |
| HG03566 | MSL | HG03547 | 0.364 | HG03454 | 0.329 |
| NA19150 | YRI | HG02325 | 0.363 | NA20361 | 0.347 |
| HG03033 | GWD | HG02861 | 0.342 | HG03034 | 0.349 |
| HG03034 | GWD | HG02464 | 0.334 | HG03033 | 0.349 |
| HG03339 | ESN | HG03366 | 0.342 | HG03361 | 0.347 |
| HG03361 | ESN | HG03342 | 0.334 | HG03339 | 0.347 |
| HG02762 | GWD | HG02643 | 0.345 | HG03373 | 0.335 |
| HG03250 | GWD | HG03246 | 0.345 | HG03249 | 0.340 |
| HG03493 | ESN | HG03511 | 0.345 | HG03373 | 0.335 |
| HG03373 | ESN | NA19130 | 0.344 | NA19150 | 0.342 |
| HG03306 | ESN | HG03514 | 0.344 | HG03307 | 0.332 |
| HG03307 | ESN | HG03398 | 0.343 | NA19150 | 0.342 |
| HG03312 | ESN | NA19147 | 0.343 | HG03249 | 0.336 |
| HG03309 | ESN | HG03193 | 0.343 | NA18487 | 0.341 |
| HG03249 | GWD | HG02678 | 0.343 | HG03250 | 0.340 |
| NA18487 | YRI | NA18934 | 0.339 | HG03309 | 0.341 |
| HG02869 | GWD | HG02896 | 0.341 | NA20361 | 0.336 |
| HG02965 | ESN | HG02981 | 0.341 | HG03249 | 0.335 |
| HG03508 | ESN | HG03199 | 0.339 | HG02964 | 0.334 |
| HG03076 | MSL | NA19129 | 0.339 | HG03569 | 0.335 |
| HG02964 | ESN | HG03193 | 0.339 | HG03508 | 0.334 |
| HG02478 | ACB | NA18858 | 0.338 | NA19150 | 0.337 |
| HG03582 | MSL | HG03398 | 0.333 | NA19150 | 0.337 |
| HG03569 | MSL | NA18915 | 0.336 | HG03076 | 0.335 |
| HG03408 | MSL | HG03115 | 0.334 | NA19150 | 0.330 |

**Table S3.**
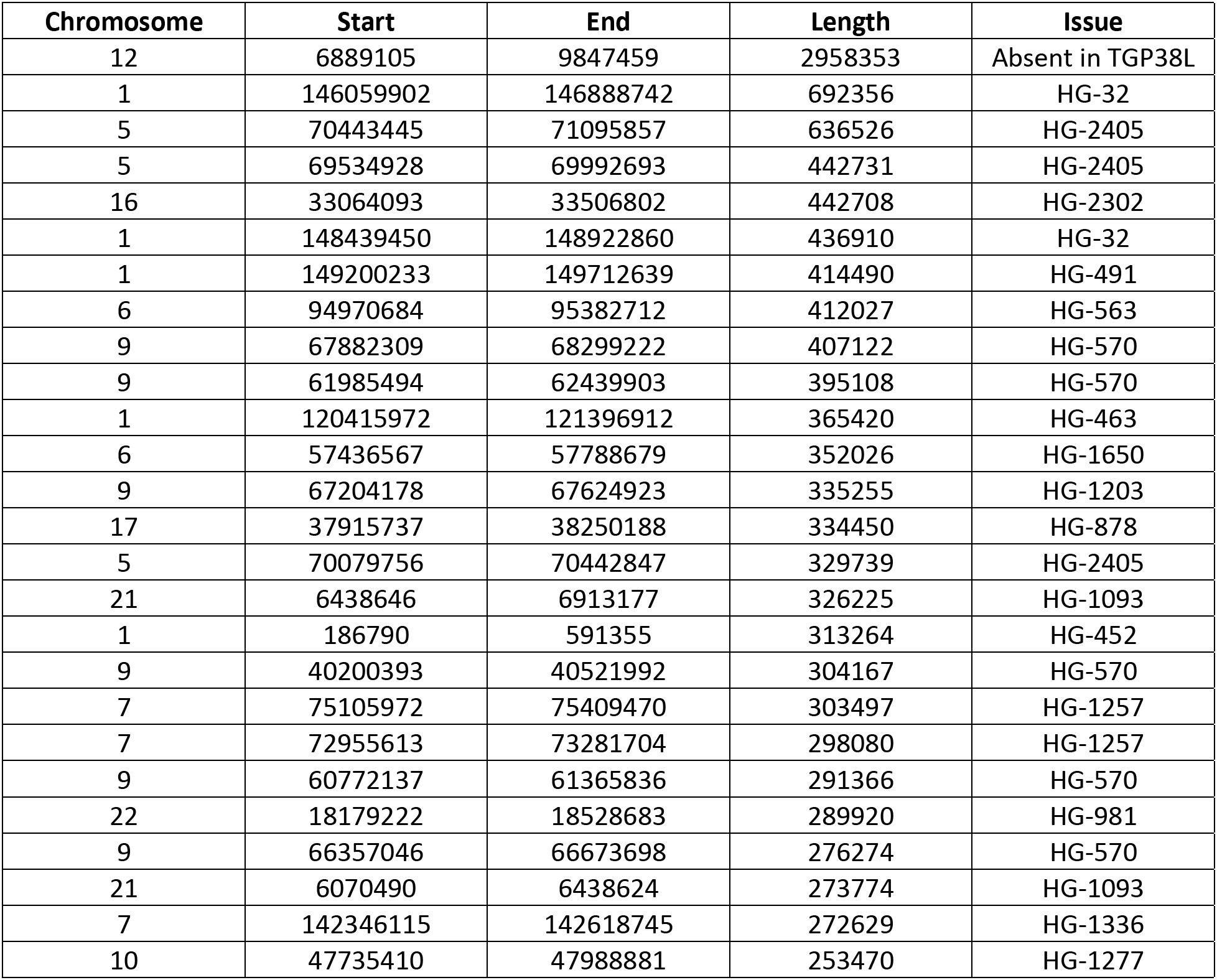
The 26 longest genomic regions enriched in discrepancies between TGP versions. Codes in the ‘Issue’ column refer to entries in the Genome Reference Consortium Incident Database (https://www.ncbi.nlm.nih.gov/grc/human/issues).

| Chromosome | Start | End | Length | Issue |
| --- | --- | --- | --- | --- |
| 12 | 6889105 | 9847459 | 2958353 | Absent in TGP38L |
| 1 | 146059902 | 146888742 | 692356 | HG-32 |
| 5 | 70443445 | 71095857 | 636526 | HG-2405 |
| 5 | 69534928 | 69992693 | 442731 | HG-2405 |
| 16 | 33064093 | 33506802 | 442708 | HG-2302 |
| 1 | 148439450 | 148922860 | 436910 | HG-32 |
| 1 | 149200233 | 149712639 | 414490 | HG-491 |
| 6 | 94970684 | 95382712 | 412027 | HG-563 |
| 9 | 67882309 | 68299222 | 407122 | HG-570 |
| 9 | 61985494 | 62439903 | 395108 | HG-570 |
| 1 | 120415972 | 121396912 | 365420 | HG-463 |
| 6 | 57436567 | 57788679 | 352026 | HG-1650 |
| 9 | 67204178 | 67624923 | 335255 | HG-1203 |
| 17 | 37915737 | 38250188 | 334450 | HG-878 |
| 5 | 70079756 | 70442847 | 329739 | HG-2405 |
| 21 | 6438646 | 6913177 | 326225 | HG-1093 |
| 1 | 186790 | 591355 | 313264 | HG-452 |
| 9 | 40200393 | 40521992 | 304167 | HG-570 |
| 7 | 75105972 | 75409470 | 303497 | HG-1257 |
| 7 | 72955613 | 73281704 | 298080 | HG-1257 |
| 9 | 60772137 | 61365836 | 291366 | HG-570 |
| 22 | 18179222 | 18528683 | 289920 | HG-981 |
| 9 | 66357046 | 66673698 | 276274 | HG-570 |
| 21 | 6070490 | 6438624 | 273774 | HG-1093 |
| 7 | 142346115 | 142618745 | 272629 | HG-1336 |
| 10 | 47735410 | 47988881 | 253470 | HG-1277 |

